# Differential expression of glycosyltransferases identified through comprehensive pan-cancer analysis

**DOI:** 10.1101/2021.06.15.448506

**Authors:** Hayley M Dingerdissen, Jeet Vora, Edmund Cauley, Amanda Bell, Charles Hadley King, Raja Mazumder

## Abstract

Despite accumulating evidence supporting a role for glycosylation in cancer progression and prognosis, the complexity of the human glycome and glycoproteome poses many challenges to understanding glycosylation-related events in cancer. In this study, a multifaceted genomics approach was applied to analyze the impact of differential expression of glycosyltransferases (GTs) in 16 cancers. An enzyme list was compiled and curated from numerous resources to create a consensus set of GTs. Resulting enzymes were analyzed for differential expression in cancer, and findings were integrated with experimental evidence from other analyses, including: similarity of healthy expression patterns across orthologous genes, miRNA expression, automatically-mined literature, curation of known cancer biomarkers, N-glycosylation impact, and survival analysis. The resulting list of GTs comprises 222 human enzymes based on annotations from five databases, 84 of which were differentially expressed in more than five cancers, and 14 of which were observed with the same direction of expression change across all implicated cancers. 25 high-value GT candidates were identified by cross-referencing multimodal analysis results, including *PYGM*, *FUT6* and additional fucosyltransferases, several UDP-glucuronosyltransferases, and others, and are suggested for prioritization in future cancer biomarker studies. Relevant findings are available through OncoMX at https://data.oncomx.org, and the overarching pipeline can be used as a framework for similarly analysis across diverse evidence types in cancer. This work is expected to improve the understanding of glycosylation in cancer by transparently defining the space of glycosyltransferase enzymes and harmonizing variable experimental data to enable improved generation of data-driven cancer biomarker hypotheses.

## Introduction

More than 600,000 deaths were attributed to an estimated 1.7 million new cancer cases in the United States in 2019 (Siegel et al. 2019), following reports of over 18 million cases and 9 million deaths worldwide in 2018. Defined as a disease characterized by abnormal cell division and invasive potential, the mechanisms by which cancer initiates and progresses can be classified into ten categories of physiologic alterations and enabling characteristics that determine cancer growth (Hanahan and Weinberg 2000; Hanahan and Weinberg 2011). Although not defined as one of the hallmarks of cancer, glycosylation and changes thereof can facilitate acquisition of these capabilities that evade innate cancer defense mechanisms and promote the tumor microenvironment, enabling cancer cells to proliferate.

Glycosylation is the enzyme-catalyzed covalent attachment of a carbohydrate to a protein, lipid, another carbohydrate, or other organic compound (Varki 2017). The resulting molecules, known as glycoconjugates, are organized into classes according to the specific molecule attached to the sugar. Glycoconjugates found in higher animals include proteoglycans, glycosphingolipids, phosphatidylinositol-linked glycophospholipid anchors, and glycoproteins (Varki 2009). Although other forms of glycosylation are discussed throughout this work, protein glycosylation is a post-translational modification whereby a sugar or assembly of sugars, called a glycan, is most commonly covalently attached to a nitrogen atom of asparagine (Asn) in N-linked glycosylation, or an oxygen atom of serine (Ser) or threonine (Thr) in O-linked glycosylation in a polypeptide chain (Mueller and Meador-Woodruff 2020). For all types of glycoconjugates, additions and cleavages of individual monosaccharide units are catalyzed by a specific enzyme, or a specific set of enzymes, known as the glycosyltransferases (GTs).

Glycosyltransferases are responsible for the extension of glycan chains, and glycosidases or glycosyl hydrolases are responsible for removing sugars to form intermediates that are subsequently extended or branched by the GTs (Varki 2017). Although additional enzymes exist which are responsible for other glycan modifications like sulfation and phosphorylation, the glycosyltransferases represent the largest group of enzymes involved in glycosylation, comprising up to 2% of genes in the human genome (Varki 2017). Most individual GTs catalyze specific reactions such that one GT will not catalyze different linkages and may be the only enzyme to catalyze a given linkage. However, there are cases in which more than one GT may catalyze the same linkage, and cases in which an individual enzyme can catalyze multiple linkages (Varki 2017). In contrast to glycosyl hydrolases, there are relatively few GT inhibitors (Varki 2017), making the GTs potentially formidable contributors to disease. GTs can be specific to a given type of glycosylation, for example, O-GalNAc glycosylation, but many GTs are glycosylation pathway nonspecific, contributing to the synthesis of both N- and O-glycans, as well as glycolipids (Varki 2009; Joshi et al. 2018).

While some individual human GTs have been well-characterized, the definition of the set of GTs has historically been variable in literature. Some previous efforts to define glycosyltransferases have been stringent and limited to only those enzymes with well-known structure and experimental evidence for function, such as the subset of GTs comprising the Consortium for Functional Glycomics (CFG) (CFG 2020) array. Other attempts have been more inclusive, such as the systematic classification of enzymes into families based on sequence similarity as conducted by the Carbohydrate Active Enzymes Database (CAZy) (Lombard et al. 2014). Combining these classifications with functional annotations assigned to enzymes as part of curation efforts through Gene Ontology molecular function terms (Ashburner et al. 2000; The Gene Ontology 2019), InterPro families (Mitchell et al. 2015), and Pfam domains (Finn et al. 2016) could better define the space of GTs by annotating enzymes having shared features with known GTs. This process would augment the available evidence for an enzyme’s role as a GT and, by applying an additional layer of curation and review, result in a complete and well-supported list of human glycosyltransferases. The generation of such a list would represent a comprehensive, evidence-supported, transparent, and reproducible process, culminating in a valuable product for community consumption.

Despite the lack of uniform definition of the GT space, there is considerable literature evidence supporting the role of human GTs in disease. In fact, many individual GTs have been reported with diverse disease associations, including but not limited to non-alcoholic fatty liver disease (NAFLD) (Zhan et al. 2016), familial amyotrophic lateral sclerosis (Cooper-Knock et al. 2019), and schizophrenia (Mueller and Meador-Woodruff 2020). A recent review by Reily et al. (Reily et al. 2019) surveyed the impact of glycosylation in human disease, reporting individual GTs with roles in congenital disorders of glycosylation (CDG), immunity and inflammation, and cancer. Moreover, individual GTs have been implicated in cancers of the skin (Christiansen et al. 2014; Tang et al. 2019), bladder (Wahby et al. 2020), breast (Christiansen et al. 2014), colon (Christiansen et al. 2014; Deschuyter et al. 2020), kidney (Drake et al. 2020), liver (Christiansen et al. 2014; Angata et al. 2020), prostate (Itkonen et al. 2020), and ovary (Christiansen et al. 2014). High mobility group box 1 (*HMGB1*) alone has been implicated in many cancers including breast, colon, lung, blood, bone, pancreas, liver, kidney, stomach, and skin, and has been observed to act as both tumor suppressor and oncogenic factor depending on its location and post-translational modifications (PTMs), including N-glycosylation (Richard et al. 2017).

In addition to studies reporting evidence for roles of individual GTs in cancer, other studies have sought to characterize the roles of entire groups of GTs involved in either a specific synthesis pathway or a specific disease. For example, fucosyltransferases involved in sialyl Lewis X (sLe^X^) synthesis were found to be over-expressed in prostate cancer (PCa) cells as compared to normal prostate tissue (Barthel et al. 2008). Similarly, enzymes involved in the glycosylation of the αDG receptor, a loss of which is associated with increased mortality in patients with Renal cell carcinoma (ccRCC), were observed to have an association between downregulation and increased overall mortality (Miller et al. 2015). mRNA expression of several GTs has been found to be significantly altered in colorectal adenomas and carcinomas, with distinct expression changes for distinct GTs(Petretti et al. 2000). O-linked mucin-type glycosylation is altered in over 90% of breast cancers, frequently categorized by increased sialylation and more linear sugar chains (Burchell et al. 2018). UDP-glucuronosyltransferase (UGT) genes have been studied and polymorphisms thereof have been identified as genetic risk factors for bladder, breast, colorectal, and other cancers (Hu et al. 2016).

The landscape of large-scale, comprehensive and systems biology type studies is constantly growing, but there are currently few comprehensive studies analyzing all GTs. The recent analysis of GTs in pancreatic cancer by Gupta et al. (2020) (Gupta et al. 2020) represents a thorough characterization of GTs in a single cancer type, and the 2016 study by Ashkani and Naidoo (Ashkani and Naidoo 2016; Gupta et al. 2020) analyzing GT expression profiles in six tumor types represents a comprehensive characterization using microarray data. In this study, we describe an approach comparable to these two studies using RNA-Seq derived data from more cancer types, presenting a survey of differentially expressed GTs (DEGTs) in individual cancers and across multiple cancer types.

While analysis of mRNA-Seq derived paired tumor and adjacent normal counts data enables identification of differential expression between tumor and normal groups, there is little direct context for interpreting the significance of a change in expression with respect to diagnosis, prognosis, or treatment: these inferences usually require overlap analysis with a secondary dataset containing functional insights. In fact, numerous other data types exist that could better inform the interpretation of identified differentially expressed genes in cancer. Comparative genomics studies represent a valuable means of informing interpretation of consequences of genetic changes through assumptions of conservation. While most related comparative genomics studies focus on comparing attributes of cancer in different organisms (Datta et al. 2005; Puente et al. 2006; Gordon et al. 2009), consideration for how the normal or healthy expression context may improve our understanding of cancer. Characterization of the interplay between microRNA (miRNA) and glycogenes has rapidly expanded in recent years (Thu and Mahal 2020), and it has been suggested that miRNA can act as a proxy to study glycogenes important to a given pathway based on this relationship (Kurcon et al. 2015). Interrogating miRNA-Seq expression data for co-occurrence of differential expression with DEGTs could better define the regulatory interactions between miRNAs and GTs, and identify mechanisms for observed differential expression of GTs. Literature on the role of glycosylation and glycogenes in cancer is dense and growing, proving difficult to search through traditional query syntax. Moreover, assertions of expression in cancer in literature are highly variable in strength and confidence, due in large part to structural variations in grammar used to communicate such findings. Automated literature mining approaches can lower barriers to systematic capture of gene-expression relationships from thousands of small scale-studies (Gupta et al. 2018), and these relationships can be used to strengthen or refute experimental findings, for example, of differential expression of GTs in cancer, toward an improved prioritization of potential biomarkers for downstream study. Thus, integration of healthy expression context, miRNA-Seq derived differential expression in cancer, and automatically mined literature about differential expression in cancer may provide better evidence by which to prioritize GTs for study as potential cancer biomarkers. Additional integration of the functional impact of possible deregulation on glycan synthesis could further identify specific glycan structures as a distinct class of biomarkers.

The goals of this study were to: (1) define a list of human glycosyltransferases and characterize their differential expression in cancers; (2) identify GT orthologs in mouse and compare differential expression in cancer with healthy expression profiles in human and mouse; (3) integrate these findings with additional experimental evidence toward the generation of high-value targets and prioritized cancer biomarker hypotheses; (4) make findings publicly available, adhering to best practices for data provenance to promote reusability of the data and processes developed herein, supporting the sustainability of the cancer genomics data ecosystem. To this end, 222 human enzymes were identified to be human GTs, 172 of which were found to be significantly differentially expressed in up to 13 cancers. 133 GTs were found to have mouse orthologs, seven of which had a similar healthy expression profile across both organisms. 293 pairs of miRNA and mRNA GT targets were identified as high-value pairs based on opposing directions of differential expression and significance of fold change, and 12 differentially expressed miRNAs were identified with more than 10 co-occurring DEGTs in cancer. Glycans were found to be affected by the top impacted residues based on the number of glycans containing each residue, number of differentially expressed GTs affecting linkages, and number of alternative enzymes capable of compensating for the impacted linkage. Evidences were compiled across experimental modes to rank genes based on the availability of cancer-relevant experimental data: six GT genes were identified with extant biomarker profiles, and five gene-cancer pairs from among those highest ranked were identified as having significantly different survival between expression groups. Major findings were made available as datasets at the OncoMX data portal [https://data.oncomx.org/] with a corresponding BioCompute Object (BCO) (Simonyan et al. 2017; Alterovitz et al. 2018; Patel et al. 2021) to communicate and track provenance details. Similarly, the curated list of human glycosyltransferases and BCO is available from the GlyGen data portal (GlyGen 2020), and some datasets have been prioritized for full-scale integration into the OncoMX database and web portal for cancer biomarkers and related evidence, available at https://www.oncomx.org/. Figure 1 shows high-level details of the pipeline developed and followed for this study. Scripts used in the data processing described throughout can be found at https://github.com/hmdinger/glycosyltransferases-in-cancer.

**Figure 1.**
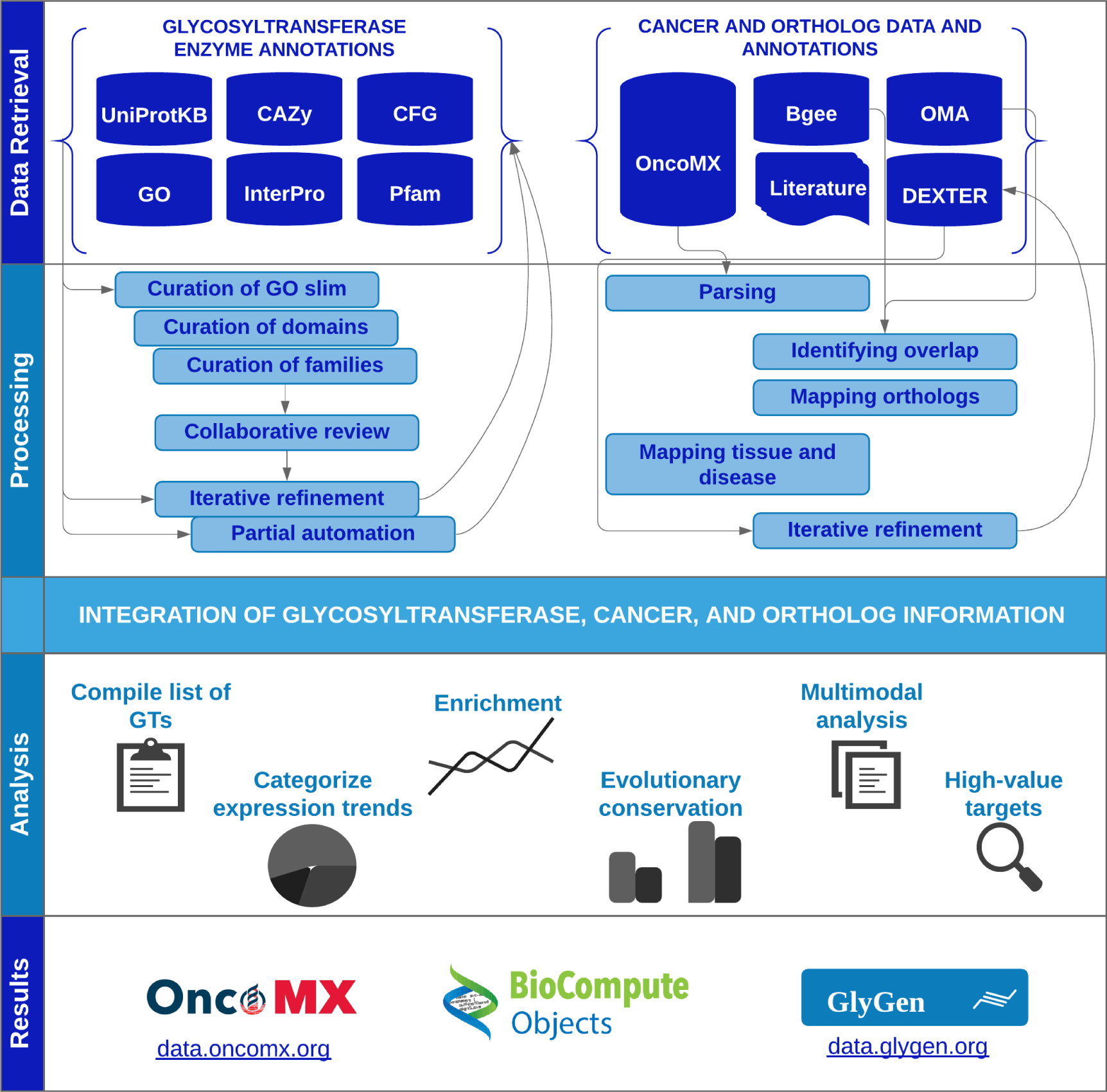
Overall pipeline. This figure shows the overarching pipeline used for generation and integration of both the curated list of human protein glycosyltransferase enzymes and supporting cancer evidence data, and the downstream analysis described herein. Annotations were retrieved from a number of resources including UniProtKB, CAZy, CFG, GO, InterPro, Pfam, and OMA (not pictured – an additional list of annotations for mouse glycosyltransferases and glycan residue annotations were retrieved from GlyGen, disease annotations from DO, anatomical entity annotations from Uberon). Experimental evidence for differential expression in cancer, “normal” assumed healthy expression, miRNA-Seq derived differential expression in cancer, and literature reporting disease involvement were retrieved from OncoMX, Bgee, and DEXTER. Preliminary data were subjected to various processing steps including manual curation and collaborative review, and generation workflows iterated over as needed. Full integration and subsequent analysis was facilitated by mapping of disease and anatomical ontology terms. Genes in the newly compiled list of curated glycosyltransferases were examined with respect to differential expression in cancer, enrichment among all cancer DEGs, conservation of normal expression profiles across human and mouse, and various additional multimodal data to identify high-value targets with respect to available cancer evidence and likelihood of functional impact. A subset of resulting datasets were made publicly available from persistent IDs in both OncoMX and GlyGen, accompanied by appropriate provenance details as defined by BCO.

The findings presented herein lay the groundwork for both a more comprehensive understanding of glycosylation in cancer, as well as an improved mechanism for identifying and prioritizing potential glycosylation biomarkers for in-depth validation studies. Moreover, the analysis framework developed for this study can be adapted to study other glycosylation related enzymes, or more generally, to study any enzyme group across diverse experimental evidence in cancer. Additionally, the variety of individual unified datasets, integrated analysis across experimental types, and availability of such information in production-level web portals represent valuable deliverables to the cancer and glycobiology research communities, including individuals in academia, industry, and the regulatory domain. We anticipate that the integration of such information into a community biomarker resource will ultimately facilitate improved cancer biomarker research.

## Results

### Identified glycosyltransferases and pipeline refinement

An early instance of the GT identification pipeline resulted in 269 proteins with evidence for roles as human GTs, 49 of which were added to the review list. Following expert review, two of these proteins were retained, and the rest excluded, with the notes pertaining to each decision maintained for future reference. The product was a curated list of 222 human glycosyltransferases (Supplemental Table S1).

Following successive rounds of feedback, the pipeline resulted in the retrieval of 284 proteins (Supplemental Table S2) with evidence for roles as human GTs based on revised automated steps of the pipeline and the static, manually curated inputs described in Methods. Upon comparison with the published list of 222 enzymes, 65 non-matching entries were flagged for review, 47 of which were already in the cumulative review list with decision to exclude. Eighteen newly identified entries were added to the review, eight of which were suggested to be retained in the next update based on available evidence. Figure 2 shows a schematic representation of the GT identification workflow. (See Supplemental Table S3 for the list of manually reviewed entries with notes and annotation for exclusion and retention in the list of human GTs.)

**Figure 2.**
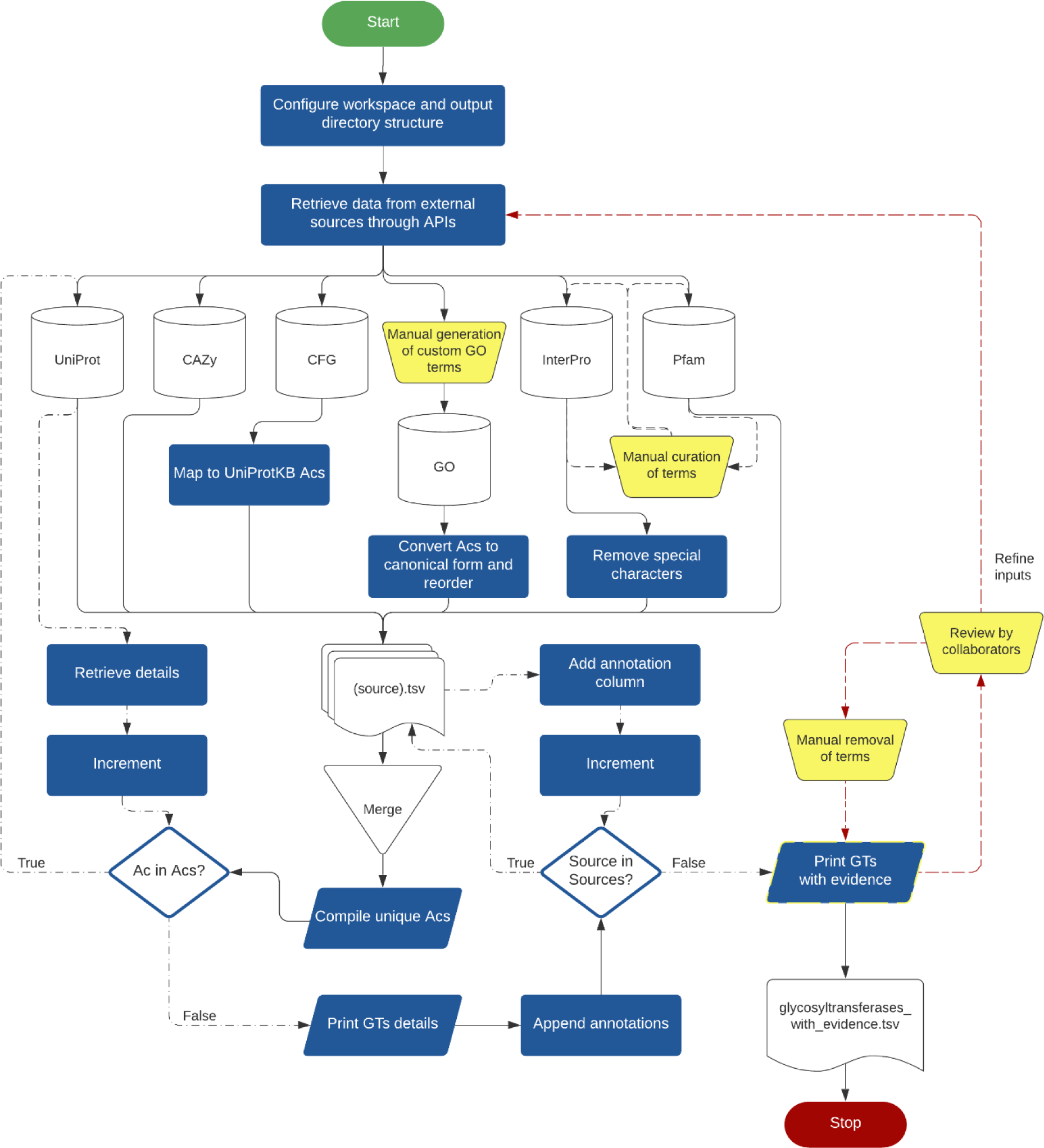
Generation of a curated list of human glycosyltransferases. This schematic representation shows all major steps of the protocol for generating the curated list of glycosyltransferases, with clear distinction of manual (yellow) and automated steps.

### GT evidence retrieved by source and considerations for curation

The list of 222 human GTs was compiled from a total of 1,074 unique evidence hits to the automated criteria (957 following manual curation and exclusion of unsuitable entries). Table 1 shows the distribution of evidences retrieved from distinct sources for automatically retrieved annotations, as well as for the curated set.

**Table 1.** Distribution of evidences for glycosyltransferases retrieved from various sources.

| Source | All hits | Curated hits |
| --- | --- | --- |
| CAZy | 203 | 195 |
| UniProt | 230 | 195 |
| CFG | 76 | 76 |
| GO | 262 | 205 |
| InterPro | 165 | 154 |
| Pfam | 138 | 132 |
| *Total* | *1074* | *957* |

Comparison of evidence across sources shows that CAZy entries have the most overlapping genes with other sources in both the total set of retrieved evidence and the curated subset. The 195 curated entries retrieved according to CAZy glycosyltransferase family annotations corresponded to 44 unique families based on sequence similarity. CFG did not contribute any unique genes, suggesting that CFG annotations were well-covered by other sources. The high gene counts contributed by GO alone, as well as the overlap of genes having GO and UniProtKB annotations, suggests a relationship between the UniProtKB kw 0328 Glycosyltransferase keyword and GO term GO:0016757 transferase activity, transferring glycosyl groups. The sharp decrease in the number of genes contributed by GO in the set of curated evidence as compared to the set of all retrieved evidence implies that the GT relevant slim contains terms not well-matched to the specific subset of molecular function under consideration. All UniProtKB hits came from a single queried keyword, and 205 GO annotations corresponded to 121 child terms under GO:0016757. Of 154 InterPro annotations and 132 Pfam annotations, 47 and 40 reduced to unique entries, respectively. The decrease in the number of genes contributed by InterPro in the subset of curated evidence as compared to all evidence implies an issue similar to that of GO terms with non-matching terms in the query set. More than 77% of identified genes have evidence from at least four sources. Figure 3 displays the overlap of evidences for GTs reported by individual contribution sources.

**Figure 3.**
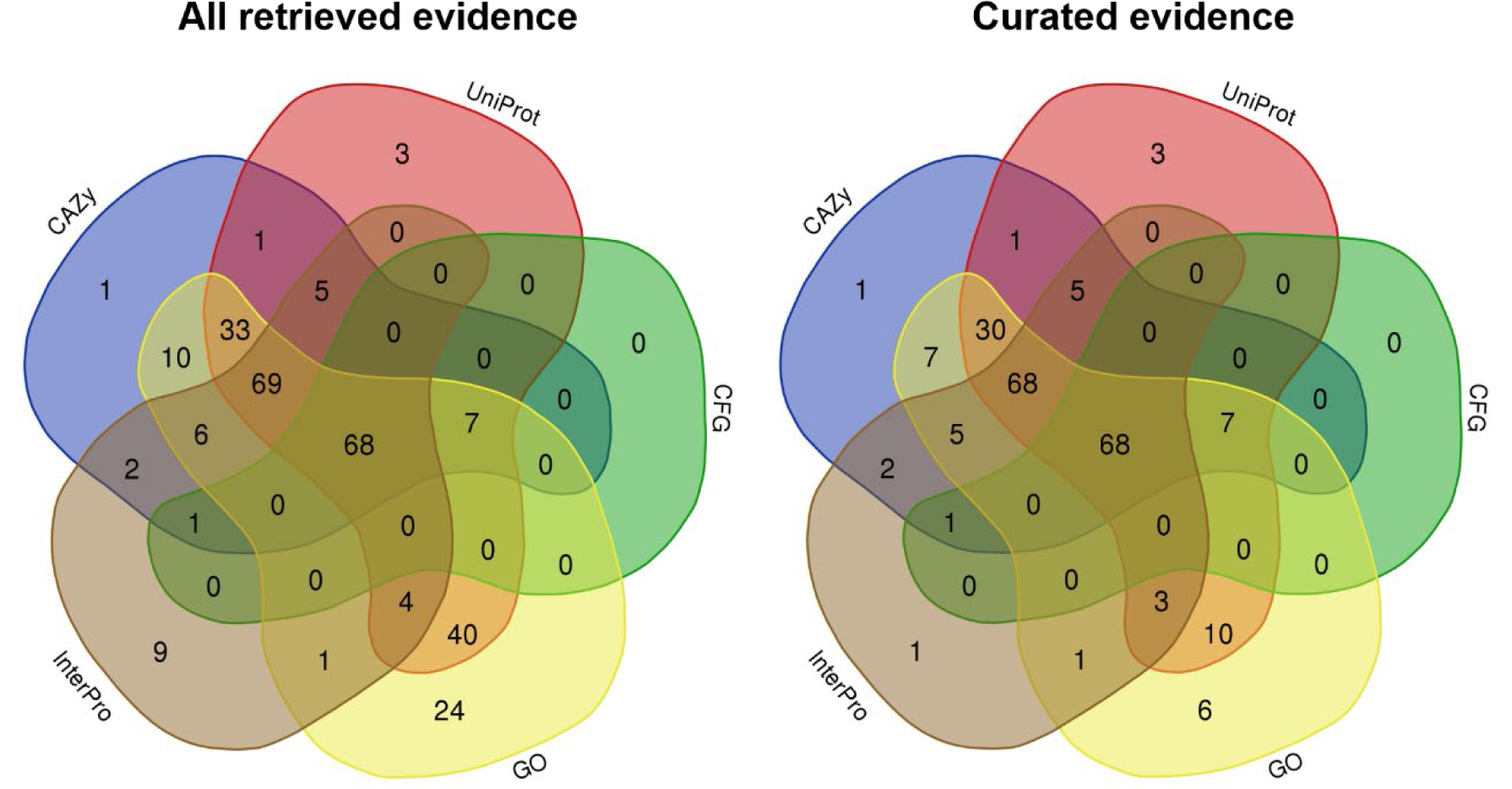
Overlap of evidence for glycosyltransferases from different sources. (Note: Pfam terms are not included as the subset of relevant terms were appended in an ad hoc fashion and not used as criteria for retrieval of glycosyltransferases.)

### Post-processing of GT dataset and availability

The resulting set of 222 human GTs and accompanying evidence was processed by the GlyGen model for dataset standardization, integration, and quality control as described by Kahsay et al. (Kahsay et al. 2020). In short, the output from the identification process described in Methods was versioned and stored in the GlyGen backend server. Additional fields including species, status, and brenda_ec_number were mapped and appended, followed by quality control (QC) steps including mapping to canonical UniProtKB accessions, mapping of field values to standard identifiers (when possible), ensuring correct formatting, and others. The processed dataset was assigned the GlyGen identifier GLY_000004 and a BCO (Alterovitz et al. 2018) was created to transparently communicate details of the workflow used in dataset generation. The processed dataset of human GTs and the corresponding BCO JSON will be maintained hereafter by the GlyGen resource (GlyGen 2020), and will be made available to all users as a dataset object at https://w3id.org/biocompute/portal/GLY_000004, or through programmatic access via API from https://api.glygen.org.

Of note, during the period of analysis described in this work, two additional GT list resources were created and published by outside research groups (referred to as external GT lists throughout) (Hansen et al. 2015; Joshi et al. 2018). These lists were not referenced prior to expression analysis of the 222 GTs defined herein, but were assessed for overlap of contained enzymes in a post-hoc fashion. Three enzymes (*AGL*, *DPAGT1*, and *NDST1*) were observed to be absent from both external lists, and, upon further manual inspection, were flagged for removal for removal from the human GTs list defined in this work (referred to as internal GT list throughout). A fourth enzyme, *MGAT4D*, was identified as belonging to only one of the external lists, and was also flagged for removal based on the finding of its primary activity as an inhibitor of *MGAT1* rather than GT activity (Huang et al. 2015a). Conversely, four enzymes (*TMTC1*, *TMTC2*, *TMTC3*, and *TMTC4*) were found in one of the external lists but not the internal list of 222 human GTs. Manual review identified these enzymes as newly discovered POMTs involved in the transfer of O-linked mannose glycans to cadherin domains, flagging them for inclusion in the internal dataset. Additionally, the Hansen and Joshi GT lists provided annotations related to GT activity that were determined to be valuable above and beyond evidence of GT activity. Annotations including GT fold type, glycosylation pathway, synthesis step, donor molecule, isoenzyme status, regulated status, cellular localization, and protein type, were manually appended to the internal GT list (Supplemental Table S4). These and other future changes to the list will be assigned a subsequent version number according to the parent GlyGen data release

### Overview of differential expression data

A total of 20,370 proteins belonging to the complete reviewed human proteome were retrieved from UniProtKB/Swiss-Prot. 222 of these proteins were identified as human GTs by the process described in Methods and retrieved from the curated file hosted at the GlyGen data portal. 14,297 genes were significantly differentially expressed as defined by *P* value adjusted for multiple testing by BH FDR procedure < 0.05 with a corresponding |log2FC| ≥ 1.0 in at least one of 16 cancer types. Supplemental Table S5 lists the cancers included in the analysis mapped to corresponding tissue and CDO slim terms. (See Supplemental Table S6 for counts of all differentially expressed genes (DEGs) at increasing log2FC thresholds for three versions of differential expression datasets analyzed as QC.)

### Differential expression of GTs in cancer

Of the 222 human GTs identified above, 172 were differentially expressed in cancer with BH-adjusted *P* < 0.05 and |log2FC| ≥ 1.0 in at least one of 16 cancer types, analyzed at the TCGA study level (Supplemental Table S7). (Supplemental Table S8 shows the number of DEGTs reported for different thresholds of log2FC.) Of the 172 DEGTs, 44 were observed to be over-expressed across all cancer types in which they were implicated, and 28 were observed to be under-expressed across all cancer types in which they were implicated (Supplemental Table S9). The remaining 100 GTs were observed to be differentially expressed in both directions across different cancer types. (Supplemental Figure S10 displays the distribution of DEGTs by number of cancers affected, and Supplemental Figure S11 shows the trend patterns of DEGTs across cancers for all explored log2FC thresholds.) A heatmap was generated for all GTs with *P* < 0.05 after adjusting for multiple testing, showing the differences across cancer types (Figure 4 panel A). Esophageal carcinoma had the fewest DEGTs with 57, and kidney renal clear cell carcinoma the most with 171, breast invasive carcinoma and lung squamous cell carcinoma tied for second at 167, and colon adenocarcinoma next with 164. The top 40 ranks were computed based on the highest absolute value of log2FC for DEGTs, resulting in 26 unique GTs differentially expressed in at least one of 12 different cancers (see Table 2). All individual GTs in this high-value subset were differentially expressed in multiple cancers.

**Figure 4.**
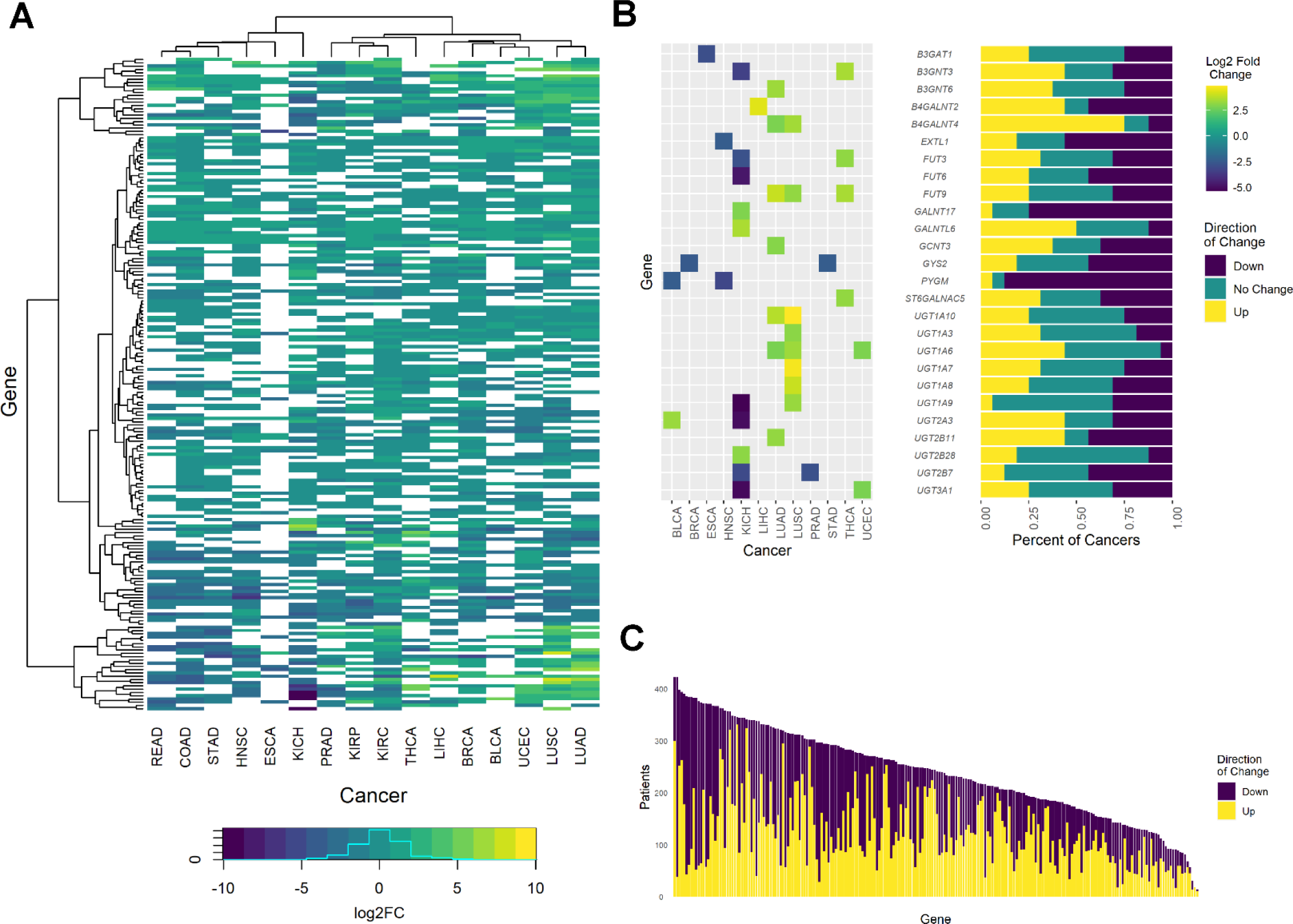
Overview of differentially expressed glycosyltransferases in cancer. A) Heatmap of P values for all DEGTs. B) Heatmap of log2FC for top 26 DEGTs ranked by magnitude of log2FC with summary of percentage of samples for that gene having pairwise increase, decrease, or no change in expression across all cancers. C) Distribution of log2FC for genes across all cancers of all patients having pairwise increase or decrease of expression.

**Table 2.**
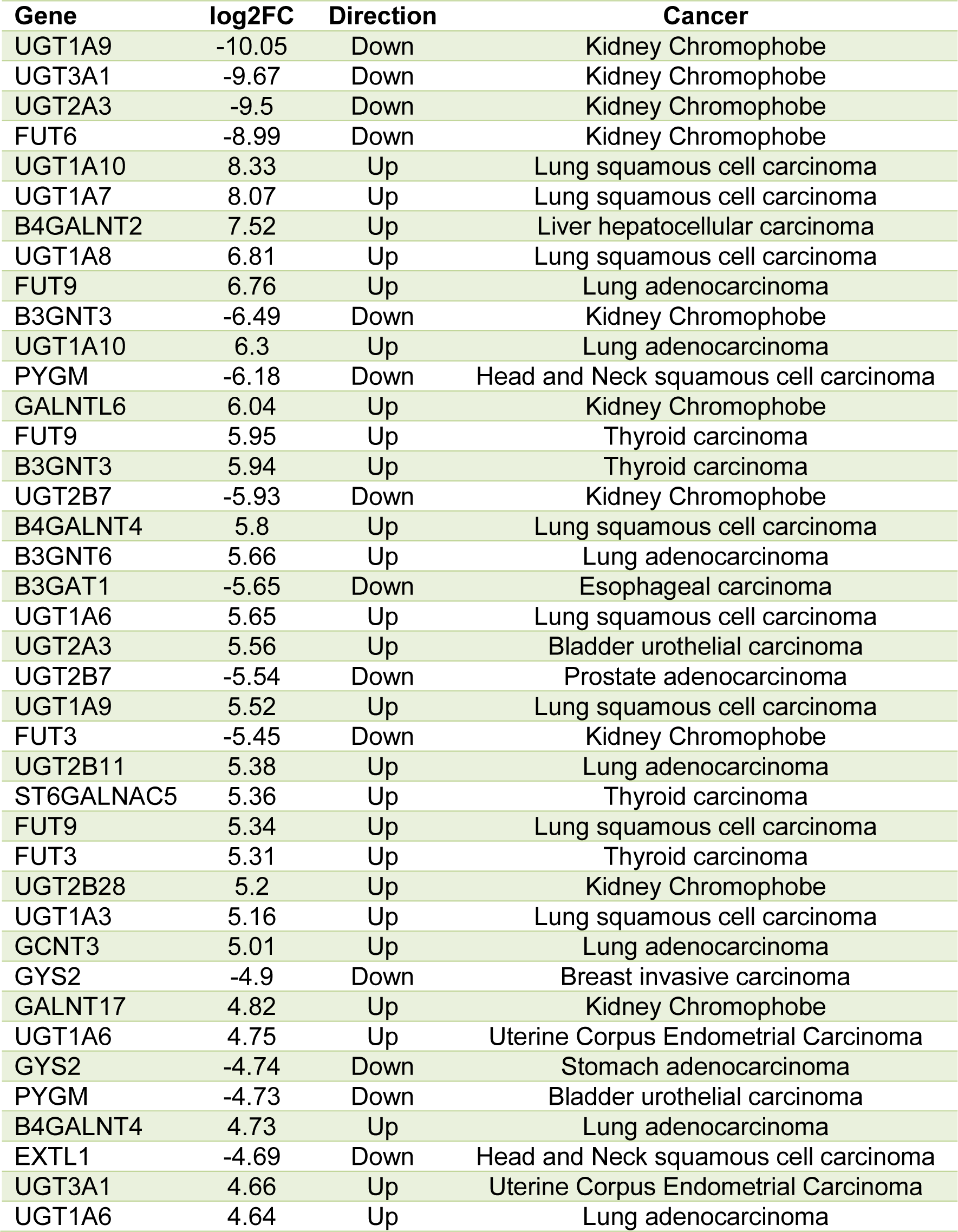
Top 40 differentially expressed glycosyltransferases by log2FC.

A second heatmap was generated for the top 26 DEGTs, and patients with paired tumor and adjacent normal samples were counted with respect to each gene to report the frequency of patients whose pairwise expression increased, decreased, or was not observed to change in tumor samples as compared to normal samples (Figure 4 panel B). *B4GALNT4* was the most over-expressed across cancers among the top ranked genes, over-expressed in 12 cancers. *PYGM* was the most under-expressed, with under-expression across 12 cancers. All genes were summarized with respect to the total number of samples having pairwise trends of up or down across all cancers and ranked in order from most sample counts to least (Figure 4 panel C). *GALNT16* and *COLGAT1* had the most samples with pairwise increased expression across cancers, and PYGM had the most samples with pairwise decreased expression across cancers. (Supplemental Table S12 shows the top genes ranked by patient counts across samples.)

*PYGM* and *B4GALNT4* had the most total samples with data, each having 422 samples. *GLT6D1* had the smallest number of total samples at 14, with 10 increased and four decreased. LALBA had the second least number of total samples, but had the least decreased samples at zero, observed to have pairwise increase in all 19 samples.

### Enrichment

Enrichment analysis determined that the DEGTs in cancer (172 of 222) were not enriched amongst the set of all cancer DEGs based on expectation calculated from the frequency of 14,297 human cancer DEGs (adjusted *P* = 1.00). GTs were only enriched among the set of differentially expressed genes in prostate adenocarcinoma (*P* = 0.047). (Supplemental Table S13 summarizes gene enrichment findings for DEGTs in cancer.)

A second analysis assessing enrichment of GT families in cancer similarly identified that there was no significant enrichment of GT families amongst the set of all DEGs in cancer. With respect to specific cancer types, GT families 1, 12, 27, and 29 were found to be over-represented in at least one cancer type, with GT1 over-represented in eight different cancer types (Supplemental Table S14). No GT families were under-represented with respect to differential expression in cancer.

### Identification of orthologs and similarity of healthy expression of GTs

165 of the 222 human GTs were reported as part of a 1:1 ortholog pair from OMA. 133 of the remaining 165 were also found to be associated with a mouse GT enzyme annotated in the GlyGen dataset (Supplemental Table S15). Only seven DEGTs were identified as high-value GTs in this analysis, having a cancer trend opposing the observed similar trend in normal expression across human and mouse orthologous genes (Table 3). Two of the seven genes were reported with two high-value similarity relationships in different cancers.

**Table 3.** High-value differentially expressed glycosyltransferases based on similar normal expression.

| Human Gene | Cancer | Trend | Log2 FC | Organ | Human | Mouse |
| --- | --- | --- | --- | --- | --- | --- |
| PYGM | Breast invasive carcinoma | Down | -1.88 | breast | HIGH | HIGH |
| PYGM | Esophageal carcinoma | Down | -3.19 | esophagus | HIGH | HIGH |
| B4GALNT3 | Kidney renal clear cell carcinoma | Down | -3.46 | kidney | HIGH | HIGH |
| ST6GALNAC2 | Kidney renal clear cell carcinoma | Down | -2.18 | kidney | HIGH | HIGH |
| ST6GALNAC2 | Prostate adenocarcinoma | Down | -1.15 | prostate gland | HIGH | HIGH |
| PYGB | Bladder urothelial carcinoma | Down | -1.32 | urinary bladder | HIGH | HIGH |
| MFNG | Lung squamous cell carcinoma | Down | -1.99 | lung | HIGH | HIGH |
| POMGNT2 | Kidney renal clear cell carcinoma | Down | -1.01 | kidney | HIGH | HIGH |
| NDST1 | Lung squamous cell carcinoma | Down | -1.03 | lung | HIGH | HIGH |

Scores were computed as described in Methods (Supplemental Table S16) and counts summarized for both the sum and average of scores across all samples in all tissues (Supplemental Figure S17). Per gene visualizations were generated for integration into OncoMX to enhance utility (see Figure 5).

**Figure 5.**
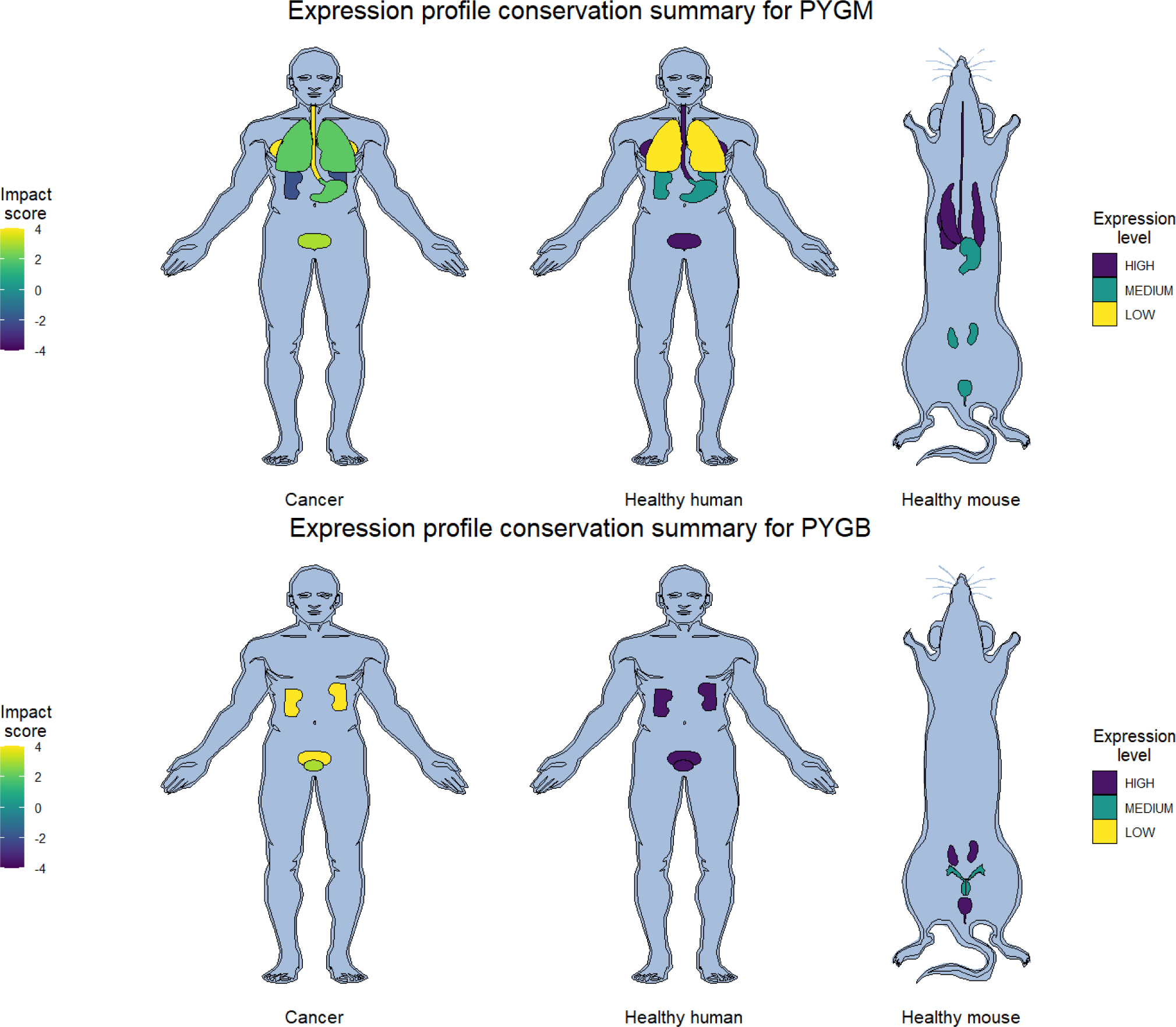
Visualization of trends of expression in cancerous human, healthy human, and healthy mouse samples.

### Differentially expressed glycosyltransferases as miRNA targets

A total of 56 miRNAs were retrieved with targets of 7,328 genes in 34,091 unique miRNA-target pairs. A subset of 37 miRNAs were found to target 54 GTs in 191 unique miRNA-GT target pairs. 530 miRNA-GT target pairs were identified with opposing expression trends, 293 of which were determined to be high-value as defined by both mRNA and miRNA log2FC magnitude of at least 1.0. The set of high-value candidates includes pairing combinations of 30 unique GTs targeted by 32 unique miRNAs in 14 different cancers. Table 4 includes the top 20 high-value miRNA-GT target pairs. Supplemental Table S14 includes all high-value miRNA-GT target pairs.

**Table 4.** Top high-value miRNA-target pairs with differential expression in cancer.

| Gene | Cancer | GT trend | GT log2FC | miRNA | miRNA trend | miRNA log2FC |
| --- | --- | --- | --- | --- | --- | --- |
| MFNG | Lung squamous cell carcinoma | Down | -2.19 | MIR210 | Up | 4.77 |
| PYGM | Lung squamous cell carcinoma | Down | -2.19 | MIR210 | Up | 4.77 |
| GALNT16 | Lung squamous cell carcinoma | Down | -2.06 | MIR210 | Up | 4.77 |
| GALNT17 | Bladder urothelial carcinoma | Down | -3.47 | MIR767 | Up | 4.39 |
| FUT5 | Uterine Corpus Endometrial Carcinoma | Up | 2.54 | MIR139 | Down | -4.08 |
| PYGM | Bladder urothelial carcinoma | Down | -4.73 | MIR210 | Up | 4.02 |
| GALNT17 | Bladder urothelial carcinoma | Down | -3.47 | MIR210 | Up | 4.02 |
| GALNT16 | Bladder urothelial carcinoma | Down | -2.04 | MIR210 | Up | 4.02 |
| ST6GAL1 | Kidney renal clear cell carcinoma | Down | -2.16 | MIR210 | Up | 3.75 |
| GALNT15 | Uterine Corpus Endometrial Carcinoma | Down | -2.95 | MIR877 | Up | 3.7 |
| GLT8D2 | Uterine Corpus Endometrial Carcinoma | Down | -2.56 | MIR877 | Up | 3.7 |
| MFNG | Lung squamous cell carcinoma | Down | -2.19 | MIR301B | Up | 3.62 |
| GALNT3 | Kidney renal clear cell carcinoma | Down | -2.66 | MIR4652 | Up | 3.43 |
| PYGM | Uterine Corpus Endometrial Carcinoma | Down | -2.85 | MIR210 | Up | 3.32 |
| GALNT17 | Uterine Corpus Endometrial Carcinoma | Down | -2.62 | MIR210 | Up | 3.32 |
| GLT8D2 | Uterine Corpus Endometrial Carcinoma | Down | -2.56 | MIR210 | Up | 3.32 |
| XYLT1 | Uterine Corpus Endometrial Carcinoma | Down | -2.2 | MIR210 | Up | 3.32 |
| GALNT15 | Uterine Corpus Endometrial Carcinoma | Down | -2.95 | MIR130B | Up | 3.27 |
| GALNT15 | Uterine Corpus Endometrial Carcinoma | Down | -2.95 | MIR937 | Up | 3.27 |
| GALNT17 | Uterine Corpus Endometrial Carcinoma | Down | -2.62 | MIR767 | Up | 3.18 |

### Literature-mining evidence for DEGTs in disease

234 literature evidences were reported for the set of GTs in human disease, and 25 evidences were reported for the set of GTs specifically in cancer. 54 unique GTs were represented in the literature evidence for all human disease in 87 diseases, and 18 unique GTs were represented in eight unique cancer types (Supplemental Table S19).

### Impact on glycans

3,866 unique glycan accessions mapped to a subset of 46 human DEGTs involved in N-linked glycosylation. Table 5 summarizes the number of implicated differentially expressed GTs, glycans, and residues associated with each cancer type. At a log2FC threshold magnitude of 1.0, 32 DEGTs were found to potentially impact 67 residues in 3,020 glycans across 16 cancer types. At a log2FC threshold magnitude of 3.0, nine DEGTs were found to potentially impact 16 residues in 1,719 glycans across 12 different cancers. Figure 6 shows a glycan schematic indicating the residues impacted by the top nine N-glycan DEGTs in cancer, and Figure 7 contains a flow diagram for the same set of DEGTs.

**Table 5.** Glycan impact.

| **Gene** | **# glycans** | **# residues** | **# cancers** |
| --- | --- | --- | --- |
| B4GALT3 | 1950 | 11 | 3 |
| B4GALT2 | 1950 | 11 | 2 |
| B4GALT1 | 1950 | 11 | 1 |
| FUT8 | 1811 | 1 | 3 |
| MGAT4C | 1067 | 2 | 11 |
| MGAT4A | 997 | 1 | 5 |
| MGAT4B | 997 | 1 | 3 |
| MGAT5 | 752 | 1 | 1 |
| MGAT5B | 752 | 1 | 7 |
| ST3GAL6 | 633 | 14 | 5 |
| ST3GAL3 | 633 | 14 | 4 |
| ST3GAL4 | 633 | 14 | 6 |
| ST6GAL2 | 498 | 9 | 6 |
| ST6GAL1 | 498 | 9 | 2 |
| MGAT3 | 351 | 1 | 8 |
| B4GALNT3 | 298 | 4 | 9 |
| B4GALNT4 | 298 | 4 | 13 |
| ALG3 | 266 | 2 | 8 |
| B3GNT8 | 174 | 6 | 9 |
| B3GNT4 | 174 | 6 | 13 |
| FUT6 | 170 | 3 | 9 |
| FUT9 | 170 | 3 | 9 |
| FUT5 | 170 | 3 | 4 |
| FUT3 | 170 | 3 | 9 |
| ABO | 108 | 4 | 5 |
| FUT11 | 105 | 1 | 1 |
| FUT2 | 93 | 3 | 9 |
| FUT1 | 93 | 3 | 5 |
| B3GALT5 | 66 | 3 | 9 |
| B3GALT2 | 66 | 3 | 8 |
| ALG9 | 43 | 1 | 1 |
| ALG10 | 11 | 1 | 1 |

**Figure 6.**
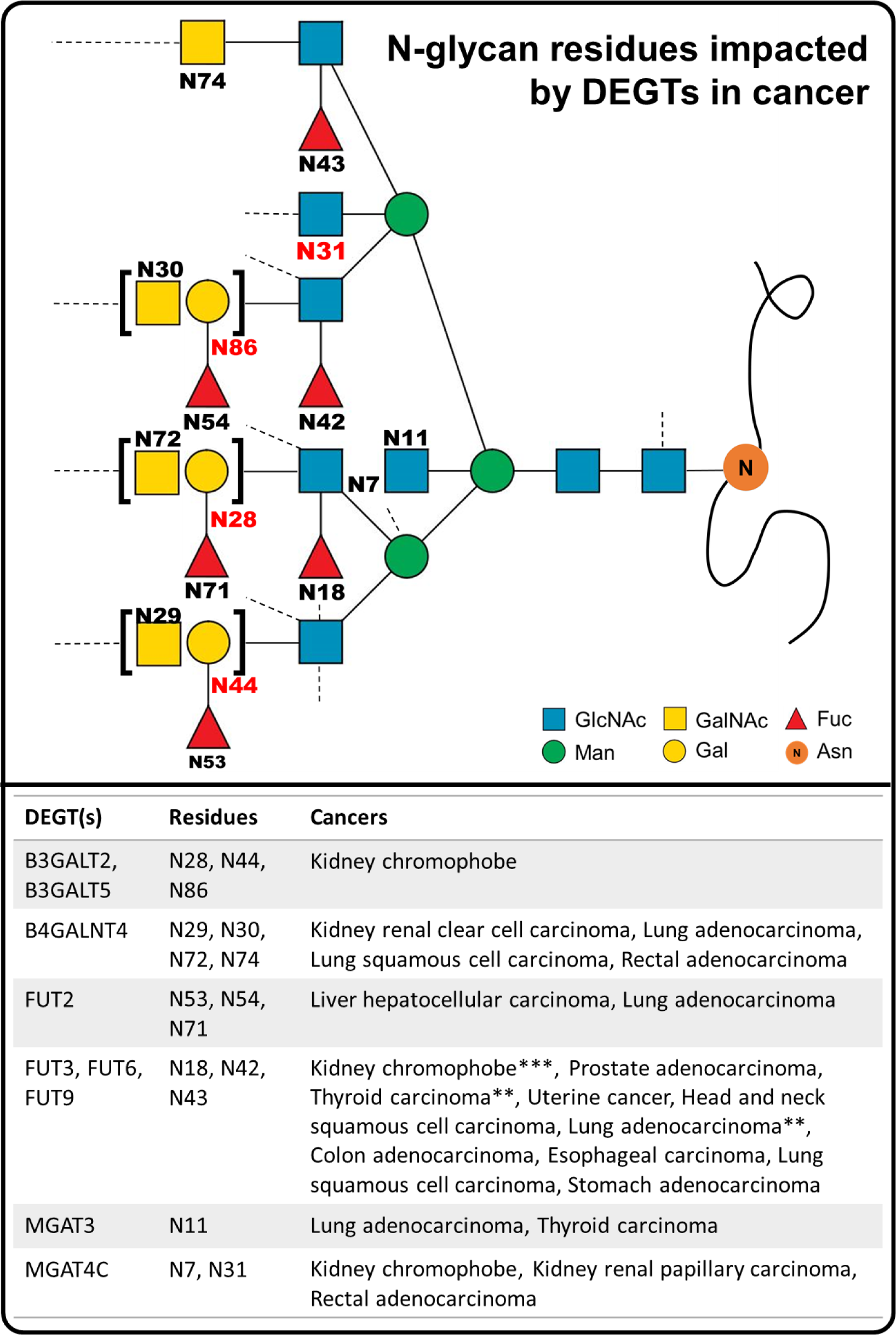
N-glycan residues impacted by DEGTs in cancer. This schematic highlights the residues that may be affected by differential expression of GTs due to the linkages catalyzed by DEGTs with |log2FC| > 3.0. Residues highlighted in red are involved in linkages for which all GTs capable of catalyzing the linkage are in the set of DEGTs. NOTE: this schematic represents thousands of combinations of residues resulting in different glycan structures. Actual glycans may span only a small subset of the residues displayed or extend beyond the implicated residue emphasized here. Brackets indicate positions where different monosaccharide residues could be linked to the same parent. Dotted lines suggest the potential for other linkages on the same residue that were not affected by DEGTs.

**Figure 7.**
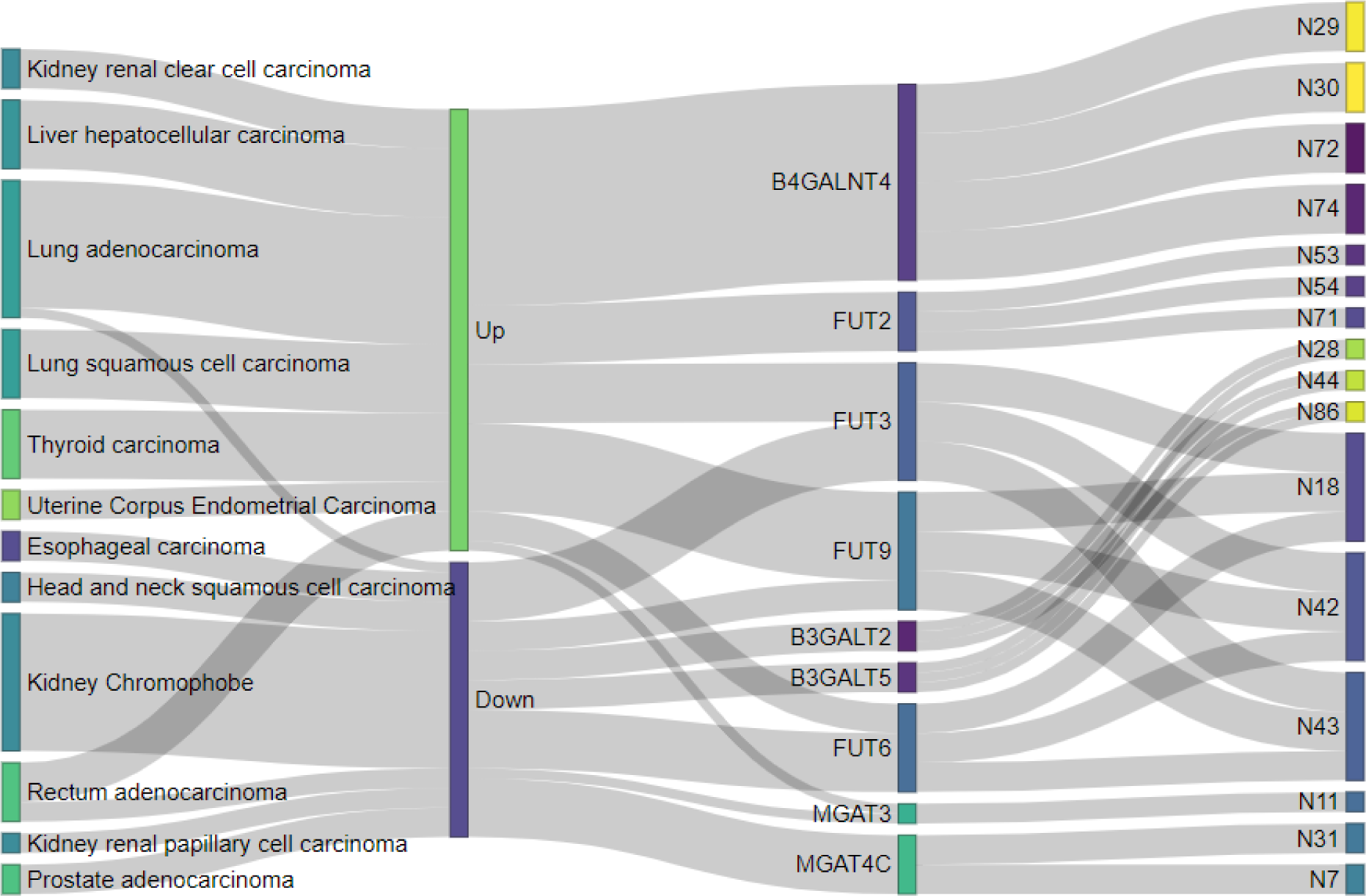
Top impacted glycan residues by DEGTs in cancer. Details for associated cancer, direction of differential expression, and specific residues impacted are displayed for DEGTs with |log2FC| ≥ 3.0.

### Overlap with existing biomarkers

Six GTs were found in the set of 939 unique EDRN biomarkers, including *CSGALNACT1*, *GALNT7*, *ALG10*, *GALNT3*, *OGT*, and *STT3A*. All but *STT3A* were also differentially expressed in cancer. A single GT, *EXT1*, was found across all datasets containing FDA approved biomarkers for various cancers. The associated organ for all of the EDRN GT biomarkers was prostate, except for *ALG10*, which was associated with breast. However, none of these genes were reported as DEGTs in prostate cancer in the DE analysis above. Similarly, the single overlapping FDA biomarker, *EXT1*, was associated with breast, but was not reported as a DEGT for breast cancer in the DE analysis described above. Supplemental Table S20 contains the merged DE and biomarker information for the DEGTs in the biomarker datasets.

### Survival analysis of high-value DEGTs

39 gene-cancer pairs (25 genes in 11 different cancers) were selected for survival analysis from the top DEGTs and miRNA targets analysis. Of the 39 total pairs, five were identified as having significantly different survival between high and low expression groups (Figure 8). Supplemental Table S21 includes a summary of findings for all survival analyses.

**Figure 8.**
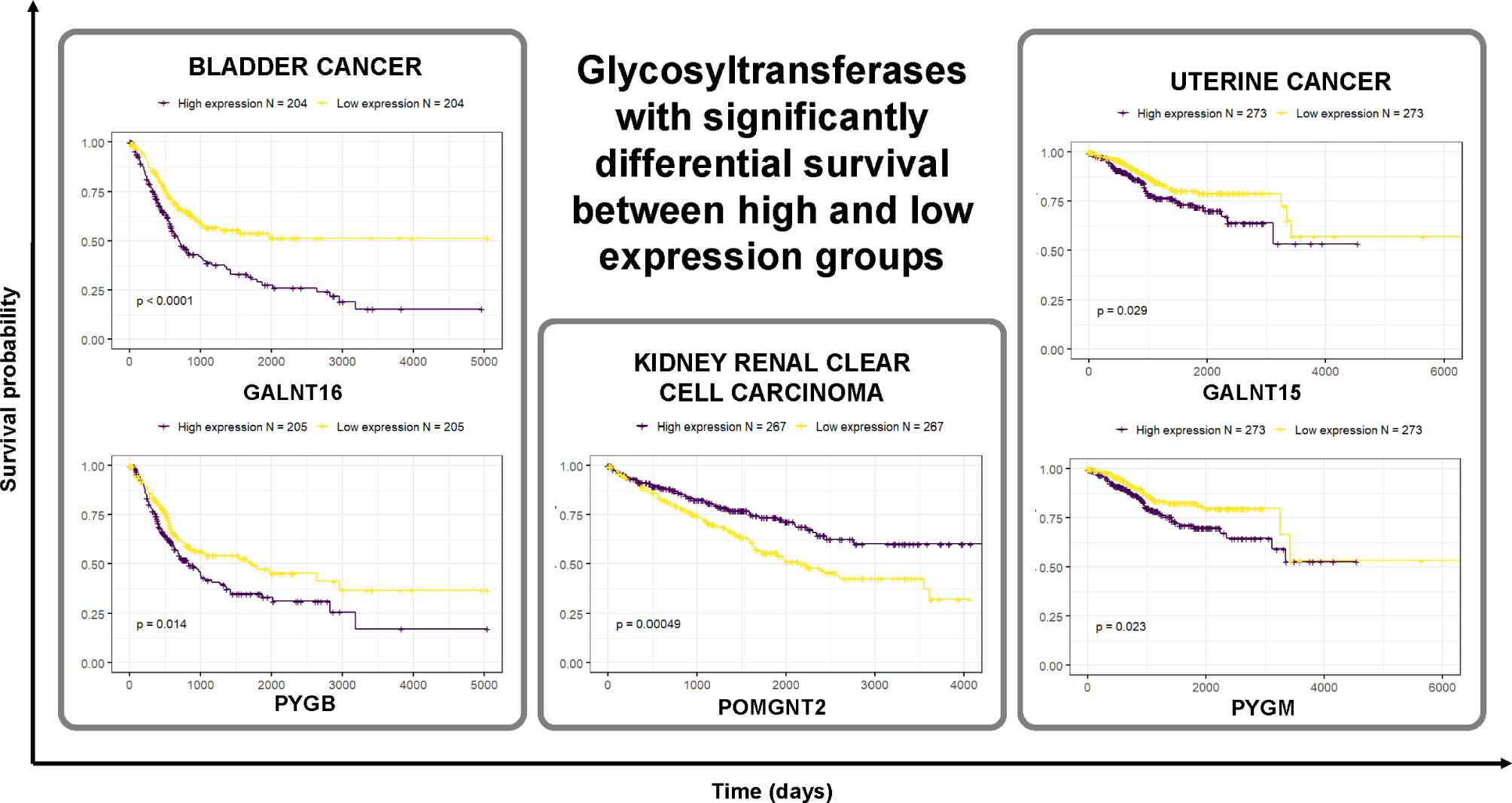
Survival analysis of high-value DEGTs in cancer. Kaplan-Meier plots of five GTs show differential survival between high expression and low expression groups.

## Discussion

### Identifying, curating, hosting, and maintaining a comprehensive list of human glycosyltransferases

The initial proof of concept for GT list generation was a largely manual undertaking.

Collaborative review identified the presence of enzymes with function not related to glycosylation, as well as terms for other processes related to sugar metabolism but not involved in glycan synthesis or attachment. Following discussion with a protein content manager for GlyGen (York et al. 2020), a second manual iteration was initiated, curating the list of child terms under GO:0016757 and flagging uncertain entries for further discussion. Flagged proteins were reviewed with respect to annotations and existing literature evidence, assigned a tentative decision, and discussed with subject matter expert before final decision of inclusion or exclusion was made. The final list of GTs comprising non-flagged and flagged but accepted enzymes was frozen and subjected to processing steps described in Kahsay et al. (2020) (Kahsay et al. 2020).

Of note, the GT generation pipeline has been partially automated following the initial release to streamline future update and review. Proteins with CAZy and UniProtKB keyword 0328 were retrieved through UniProt API. CFG was not cross-referenced in UniProt and did not have a direct export mechanism, so simple web-scraping scripts were developed to: retrieve identifiers from the Molecule page Query by Multiple Criteria results html; for each identifier, open the corresponding viewGlycoEnzyme page; and print the reference accession from the Reference tab html. GO terms from the custom slim were retrieved through QuickGO, and InterPro accessions matching search of “glycosyltransferase” were retrieved through EBI API. A master list was compiled and additional protein details including gene and protein names and EC number were retrieved from UniProt API. Pfam annotations were retrieved through EBI API for all proteins in the master list. The resulting list of annotations was manually reviewed for glycosyltransferase relevance, and remaining annotations were appended to the master list to generate the final list of human GTs.

Remaining curation steps include reviewing the list of Pfam annotations and review of the final resulting file. Although not yet automated, planned comparison of these two outputs with previous versions as well as the annotation notes and flagged list will readily identify any entries that have not yet been reviewed, reducing the curation space to a scale of ones to tens rather than hundreds. Curation is critical for high quality knowledgebases, especially when leveraging large ontologies or resources with a global scope not custom-geared for a specific research niche. Using ontology terms enables accurate and confident annotation for downstream research and comparison, but depending on the exact task and how the ontology hierarchy is structured, a proper slim for defining some group will likely necessitate manual review of some subset of child terms. Although time consuming, dedicated curation ensures the robustness of the GT list for future consumption.

Because UniProtKB keyword 0328 is heavily overlapped by GO:0016757, this step may negate the function of the custom GO slim, and is therefore being considered for removal from future versions. Although not a computationally expensive process, existing scripts have not been optimized for performance and would benefit from defining reusable functions and reducing line counts.

As stated above, GlyGen will be maintaining the GT list as a publicly available dataset at https://data.glygen.org. Planned updates to the list will remove any enzymes not involved in the biosynthesis of glycoproteins, glycolipids, proteoglycans, glycosaminoglycans, and glycogen. As a result, genes that are regulatory and accessory subunits of GTs, such as members of the oligosaccharyl transferase (OST) and dolichol-phosphate mannose (DPM) synthase complexes, would be excluded from future versions of the list.

Genes belonging to the UDP-glucuronosyltransferase (UGT) class of enzymes will also be excluded as they are involved in detoxification of drugs and compounds but not in glycan synthesis.

Additionally, weakly annotated enzymes, such as *GGTA1P*, *MGAT4D*, and *DPY19L2P1*, which do not have any conclusive literature evidence for being a GT, will be excluded from the list. Enzymes in this category have been annotated as products of pseudogenes, probable or putative glycosyltransferase, or inactive.

Furthermore, enzymes such as *B3GNTL1*, *GLT8D1*, and *GTDC1*, which do not have literature evidence but have been classified as GT by CAZy, will be sent to a glycoenzyme expert for further review. The inclusion and exclusion of such GTs in the list will be based on the expert reviewer’s recommendation, along with the existence of new literature evidence as it becomes available. For example, *C1GALT1C1*, a chaperone belonging GT31 family in CAZy, is currently considered a GT, but recent literature evidence has found no direct glycosyltransferase activity (Ju and Cummings 2002). Thus, *C1GALT1C1* will be removed from future list versions. Enzymes like *FKRP* and *FKTN* that have been denoted as GTs in Joshi et al.(Joshi et al. 2018) will also be excluded as they are actually ribitol-5-phosphate (ribitol-5P) transferases (Cataldi et al. 2020) and not glycosyltransferases.

Development of a new data model will allow the inclusion of chaperones, non-catalytic subunits, and other enzyme classes that play an important role in glycosylation and may also be implicated in disease. As these and other proteins are identified, they will be added to the interactive GlyGen Sandbox tool [https://glygen.ccrc.uga.edu/sandbox/], a tool that provides structural representation framework for glycans that facilitates their semantic annotation with information about the relationships between their structures, biosynthesis, and biophysical properties. In this way, GlyGen will be the first resource dedicated to providing the curated and comprehensive list of GT, readily available for researchers to access, view, and download. Access to this list will not only allow be of value to expert glycobiologists, but will benefit a broader audience of biologists interested in studying a diverse array of biomedical applications.

### Comparison to existing glycosyltransferase references

Many extant studies focusing on glycosyltransferases report highly variable lists of enzymes (Gupta et al. 2020), and in some cases do not readily explain the content or origins of the enzyme list (Ashkani and Naidoo 2016). For example, Gupta et al. (Gupta et al. 2020) report a number of 207 enzymes as the list of “all known glycosyltransferases,” but the cited article (Shimma et al. 2006) reports a figure of “more than 200.” Following this citation chain to a third article (Narimatsu 2004), it was reported that there were “more than 160” human GTs cloned at the time of publication in 2004. This statement was ultimately credited to the “Handbook of Glycosyltransferases and Related Genes,” the second version of which was published in 2013, citing a figure of “almost 200” GTs that had been identified at the time, 160 of which were included in the book (Angata 2002). The list produced herein represents a current, comprehensive compilation of GTs, enabling researchers studying them to reference the set of enzymes with concrete assertions about the scope of evidence used to define them. Usage of clearer descriptive language around scientific concepts ultimately improves the transparency and accuracy of downstream studies attempting to build on prior knowledge and helps to avoid fallacies of inequivalent comparisons (2016). This is especially important for amenability to use by automated text mining and other natural language processing applications whose extracted relationships are only as good as the language of the assertions used to describe them.

Of note, since the onset of this project, Hansen et al. (Hansen et al. 2015) have published a well-characterized assembly of the human glycosyltransferase genome largely defined by CAZy, sequence homology, and putative activity. Merits of their approach include addition of chromosomal and sequence attributes and fold types. The more recent update by Joshi et al. (Joshi et al. 2018) is a very comprehensive list including glycosylation pathways, a major benefit to the research community. Unique strengths of the approach described in this work include references to additional sources of evidence, primarily ontologies, and the hosting of the versioned, digitized dataset. As stated above, future updates to the list presented here will take new literature and other forms of evidence into consideration, as well as cross-referencing the lists by both Joshi et al. and Hansen et al., with special consideration regarding whether cross-reference as an additional evidence call or a more detailed integration should occur. Improvements to the organization of the list provided by Joshi et al. could drastically improve the overall utility by using machine-friendly formatting practices.

As an initial step toward integration of the Joshi and Hansen annotations into the internal GT list, genes in each of the three lists were compared, and annotations from each external work were manually appended to the corresponding GT in the internally generated list. Upon comparison, 24 genes were found in the internally generated list that were either found in only one of the external lists (eight) or absent altogether from the external lists (16). Subsequent review of these entries identified four genes for immediate removal from the GT list, including *NDST1*, *AGL*, *UGT2A1*, and *MGAT4D*, with an additional 16 under consideration for future exclusion. Conversely, 14 genes were identified from either one (seven) or both (seven) of the external lists as not currently belonging to the internally generated GT list. Four of these genes reported by Joshi et al. were flagged for inclusion: *TMCT1*, *TMCT2*, *TMCT3*, and *TMCT4* are newly discovered POMT-directed O-mannosyl-transferases. The extended table combining the evidence annotations from the approach described above and the functional annotations from the Hansen and Joshi lists can be found in Supplemental Table S4.

### Survey and enrichment of differentially expressed glycosyltransferases in cancer

When comparing the proportion of all human genes differentially expressed in at least one cancer to the total number of all human genes, a similar proportion of DEGTs to all human GTs was observed. In fact, when abundance of DEGTs in cancer were compared at different fold change thresholds, the relative proportions were comparable to those observed for all differentially expressed genes as log2FC increases (Supplemental Table S8).

When considering the direction of change among the entire pool of DEGTS, a consistently higher signal for increased expression than decreased expression was observed among glycosyltransferases. Interestingly, however, a shift towards dominant under-expression of DEGTs was observed as stricter log2FC thresholds were applied. This pattern becomes more prominent as stronger thresholds are applied with respect to *P* value, fold change, and the proportion of individual patients whose pairwise log2FC direction matches that of the trend calculated by pooling samples at the study level (Table 6). In fact, there is an almost complete shift in domination by under-expression among the GTs with high fold-change magnitude, low *P* value, and a high degree of concordance between the study level trend and pairwise log2FC trends (defined by 90% of patient pairs sharing the same trend as the study-level analysis for a given cancer).

**Table 6.**
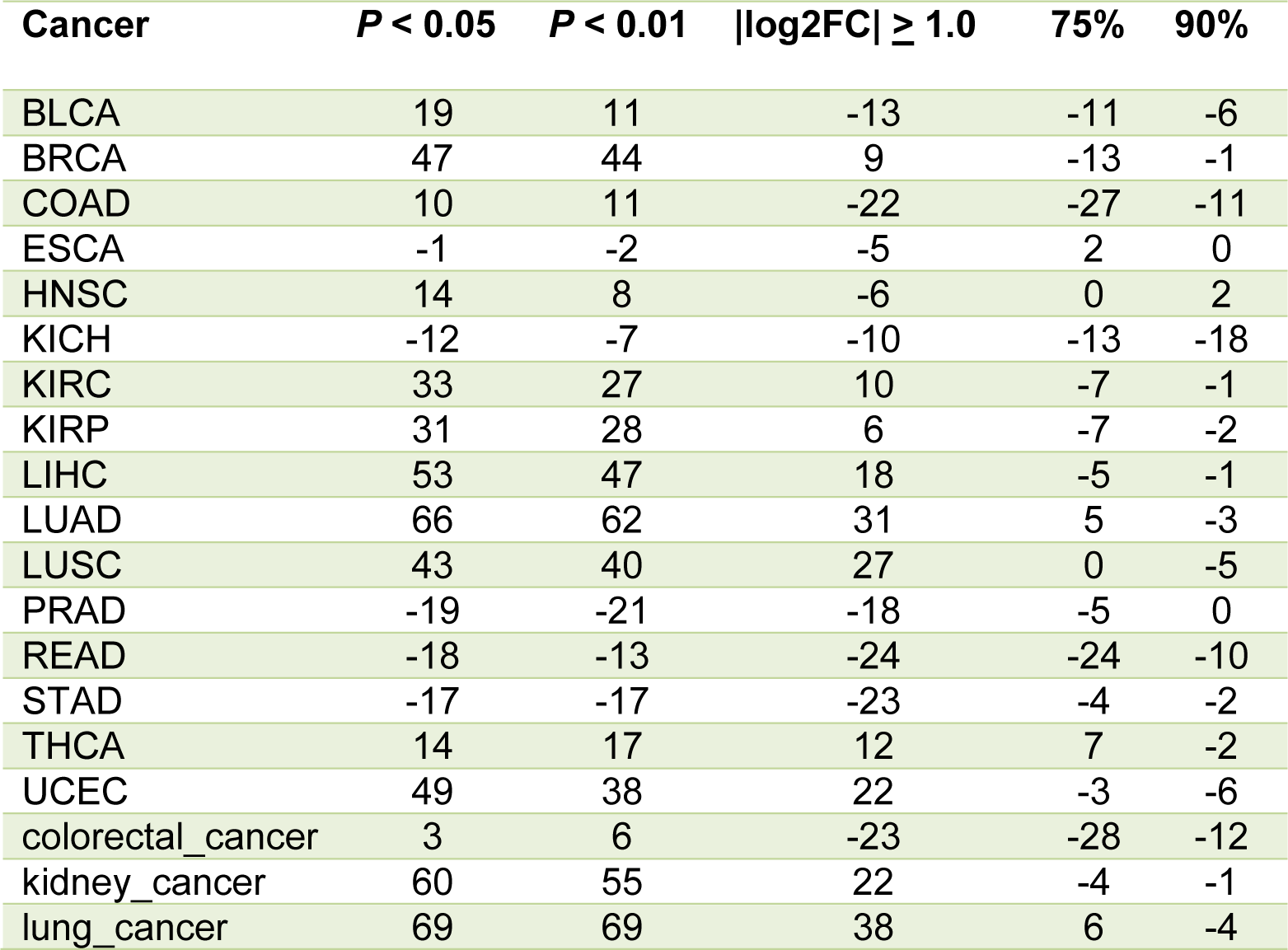
Dominating up/down trends of cancer DEGTs across significance criteria.

Rodriguez-Esteban and Jiang (2017) (Rodriguez-Esteban and Jiang 2017) reported a bias in both literature and microarray studies for over-expressed genes (as compared to under-expressed genes). Although analyzed from RNA-Seq instead of microarray, differentially expressed genes identified in this study corroborate this assertion at less stringent thresholds, but that relationship does not hold true with increasing stringency. The arguments for the bias of over-expression in literature regarding popularity of a gene and a skewed focus toward trends that are likely to be biologically meaningful are valid, underscoring the likelihood of authors to emphasize differential expression of “interesting” genes. However, the same trend observed for GTs in Table 6 holds true for the set of all reported differentially expressed genes in cancer, such that subsets of differentially expressed genes with higher fold changes are heavily biased toward under-expression. This could suggest another reason for this trend, whether it be a bias in the analysis or a true biological finding. The experimental setup used in this analysis does not lend itself to distinguishing between decreased and silent expression, but it is possible that these large size effects for decreased expression are artifacts of attempts to quantify very low abundant transcripts. Although not performed for this study, it is feasible that close investigation of the individual read counts for those samples with normal expression could verify low abundance. Summarizing the same count trends for different sets of genes, selected randomly or based on the consideration of different biological implications, could also help to determine whether this finding is specific to the GTs or ubiquitous due to some property of the analysis.

For GTs differentially expressed in multiple cancers, there are more genes with opposing expression trends than with the same trends across the implicated cancers. This can be explained in part by observations of random trials, such that with an increasing number of trials (cancers in this case), the likelihood of not observing some outcome (for example, decreased expression or increased expression for a given gene) converges to zero. Seemingly contrary to the shift in dominance of GTs with increased expression to those with decreased expression at higher levels of significance, the relative abundance of GTs whose expression trends are the same across cancers is consistently higher for those over-expressed across all cancers than those under-expressed (Supplemental Figure S11). Comparison with different disease and gene sets may help to explain the apparent incongruity between these observations.

When comparing the distribution of DEGTs across different cancer types, kidney renal clear cell carcinoma had the most with 171 and esophageal carcinoma had the fewest with 57. Literature review for implications of glycosylation and GTs in these cancers yields several reports for individual GTs or groups of enzymes in each cancer type.

Previous studies have associated increased levels of fucose and sialic acid and increased abundance of protein sugar structures with clear cell renal cell carcinoma (ccRCC) (Borzym-Kluczyk et al. 2012). Increased multi-lectin affinity chromatography (M-LAC) binding of glycoproteins and changes of plasma N-glycan levels in ccRCC suggest the presence of altered glycan structures (Gbormittah et al. 2014), and fucosyltransferases *FUT3* and *FUT6* have been observed to have decreased transcript levels in ccRCC tissue compared to matched non-tumor tissues (Drake et al. 2020). *GYLTL1B* (LARGE2) is frequently downregulated in ccRCC, associated with impaired glycosylation of dystroglycan (DG) (Miller et al. 2015).

*GALNT7* has been shown to be targeted by microRNA-214, whose decreased expression in esophageal cancer was associated with differentiation, invasion, and metastasis(Lu et al. 2016). Amplicon 3q26, containing *ST6GAL1*, is estimated to occur in more than 20% of human cancers, including esophageal cancer(Dorsett et al. 2019). *FUT3*, *FUT8*, *POFUT1*, and *POFUT2* were found to be upregulated in esophageal cancer stem-like cells (CSLC) compared to adherent cells, and *FUT8*, *POFUT1*, and *POFUT2* were expressed at higher levels in tumor tissues than matched healthy controls (Sadeghzadeh et al. 2020). Automated literature mining supplied by customized DEXTER (Gupta et al. 2018) application also identified two evidences for esophageal cancer. *ALG3* expression was higher in esophageal squamous cell carcinomas (ESCC), although the frame of comparison cannot be identified from the automated results (Shi et al. 2014). (Manual review of the publication revealed paracancerous normal tissues as the reference.) Similarly, *ST6GALNAC1* was identified to be downregulated in ESCC (Iwaya et al. 2017).

Corresponding differential expression of all of the genes mentioned above for kidney cancer are evidenced in the computed experimental results, but only a subset of the genes mentioned for esophageal cancer can be found in the computed results. Of note, there were no contradictory findings between literature reviewed and database results. Anecdotally, the log2FC size effect for each of the corroborated genes ranged from 1.19 to 2.29, suggesting that, though arbitrary, a log2FC threshold of 1.0 may be sufficient to capture true variability.

Although on a small scale, these findings serve as proof of concept that the DEGTs identified by the analysis presented herein may have biological meaning in cancer and could present valid targets for improved understanding of cancer. A large-scale comparison of literature mining results and differential expression results to better inform analysis and interpretation of differentially expressed genes in cancer is planned as part of the hardening of the OncoMX database for cancer biomarkers and related evidence (Dingerdissen et al. 2020).

### Identification and prioritization of high-value DEGTs for downstream study

Toward the identification of high-value glycosyltransferase targets in cancer, DEGTs (adjusted *P* < 0.05, |log2FC| ≥ 1.0) were ranked by absolute value of log2FC. The selection of the cutoff number for inclusion in the highest ranked genes was motivated by limiting the number of unique genes to a reasonable size for summary and downstream study. A selection of the 40 top log2FC ranked gene-cancer significant pairs resulted in a list of the top 26 unique GTs differentially expressed in at least one of 12 cancers, with magnitude of log2FC ranging from 4.64 to 10.05. Among these 26 unique genes, 11 belong to GT family 1. Despite being one of the largest GT families, this finding is consistent with the over-representation of GT1 in multiple cancer types identified through enrichment among the set of all DEGs. The overarching over-representation of glycosyltransferases belonging to GT1 in the set of all DEGs, combined with the abundant appearance among the subset of the highest ranked DEGTs, suggests that enzymes in this family could have important roles in cancer. Like GT1, most of the families in the high-value set are of the inverting type, although A and B-type folds are similarly represented. Because of the prominence of GT family 1 members, the UDP-glucuronosyltransferases, there is a heavy bias for detoxification function(Rowland et al. 2013), followed closely by an assortment of pathway non-specific enzymes responsible for capping and elongation of glycans (as annotated by Joshi et al. (Joshi et al. 2018)). UDP-glucuronosyltransferases are well known to drive the elimination of drugs, and dysregulation has been associated with the progression of various drugs (Allain et al. 2020). *UGT1A7* and *UGT1A9* may be predictors of antitumor response in colorectal cancer (CRC) patients given irinotecan and capecitabine (Carlini et al. 2005).

### High-value DEGTs with similar healthy expression profiles

Of the seven genes with identified high-value opposition between the observed similar trend of normal expression in human and mouse and observed cancer differential expression trend reported above, all seven were of the type of high normal expression and under-expression in cancer. Only one gene, *PYGM*, was previously in the high-value list as identified by ranking differential expression by magnitude of log2FC. *PYGM* has been observed to be significantly downregulated in head and neck squamous cell carcinoma (HNSCC), with decreased expression correlated with prognosis (Jin and Yang 2019). *PYGM* has also been implicated in gastric cancer through the insulin resistance pathway (Zhang et al. 2018b), and both mutation and differential expression of *PYGM* have been associated with rare breast cancers (Dieci et al. 2016). The brain isoform, *PYGB* (also implicated here), has been reported to mediate hypoxia in breast cancers, promoting metastatic phenotypes (Altemus et al. 2019).

Among the other high-value genes identified by comparing healthy and differential expression, *B4GALNT3* has been reported to be over-expressed in colon cancer (Che et al. 2014), and *MFNG* has been previously reported to be differentially expressed in gastric cancer. *ST6GALNAC2*, with computed under-expression for two cancers in the high-value expression similarity subset, has been identified as a metastasis suppressor in breast cancer, such that decreased expression results in increased metastatic burden (Murugaesu et al. 2014). *POMGNT2* is part of a prognostic panel in uveal melanoma (Luo et al. 2020), and *NDST1* is targeted and negatively regulated by miRNA-191 in glioblastoma. The abundance of literature and reported diseases for the genes fitting this high-value criteria supports the idea that evolutionarily similar expression profiles could imply conserved function and may therefore represent better targets when differentially expressed. Although outside of the scope of this project, it would be beneficial to repeat this process for different sets of genes and calculate the percent of genes with literature support meeting similar expression profile criteria.

Anatomical figures were generated to provide a visual overview of expression similarity for given tissues, with the idea that certain tissues could be more prone to similar patterns of expression across organisms than others (Supplemental Figure S17). However, no obvious association was observed between expression profile similarity and anatomical structure. Of note, feminine cancers (with the exception of breast cancer) were not included in the analysis due to sample size thresholds, so the male anatomical figure was chosen as the model for visualization. Female figures are available, however, (see Supplemental Figure S22). While these images did not reveal any great meaning for the gene set presented in this study, the theoretical relevance in communicating biological findings based on gene expression similarity exists and could be meaningful in other applications.

In addition to tissue level summaries across genes, Figure 5 clearly depicts the similarities and differences between healthy human and mouse expression values and the potential significance of the differential expression for two of the individual high-value DEGTs identified by healthy expression analysis, *PYGM* and *PYGB*. Note that the colors are intentionally offset between cancer and healthy images to highlight the exact high-value criteria captured in Table 3.

### High-value DEGTs targeted by miRNAs

miRNAs recognize response elements at the 3’ untranslated region (UTR) of their target mRNA sequences (Catalanotto et al. 2016), where they assemble into the RNA-induced silencing complex (RISC) (Macfarlane and Murphy 2010) to destabilize or prevent the translation of mRNA, ultimately silencing the gene (Catalanotto et al. 2016). The mode of silencing employed depends on the degree of complementarity in the base-pairing between the miRNA and its target sequence, such that limited base-pairing results in repression of translation and high complementarity results in mRNA cleavage (Macfarlane and Murphy 2010). Because of this negative regulatory relationship between a miRNA and its target mRNA expression, a change in expression of an mRNA would be expected to correlate with a change in expression in the opposite direction of the miRNAs targeting that mRNA.

In the study presented herein, 293 miRNA-target pairs following this pattern were identified such that the miRNA was differentially expressed in one direction in some cancer, and its reported target mRNA, that of a glycosyltransferase, was differentially expressed in the opposite direction in that same cancer. Although *PYGM* in head and neck squamous cell carcinoma is the only GT-cancer pair with observed normal expression similarity from the list of top 40 GT-cancer associations ranked by log2FC, *PYGM* appears in the list of top 20 ranked miRNA-GT pairs (Table 4) six times for five cancers (LUSC, BLCA, UCEC, BRCA, and HNSC), with two targeting miRNAs having significant differential expression in uterine cancer. Similarly, while the exact gene-cancer pair is not in the top 40 log2FC associations, several genes represented in that list are also present in the top 20 miRNA-GT pairs ranked by miRNA log2FC, including *GALNT17* (two cancers, BLCA with two miRNAs and UCEC with three miRNAs), and *FUT6* and *B3GNT6* with one miRNA each in HNSC.

miRNAs have been reportedly differentially expressed in a number of cancers, including but not limited to colorectal, lung, glioblastoma, lymphomas, and other cancers (Volinia et al. 2006). Kurcon et al.(Kurcon et al. 2015) have previously identified a set of glycosylation enzymes acting as regulatory elements through the miRNA-200 family (miR-200f) as a proxy to regulating epithelial-to-mesenchymal transition (EMT). Criteria used to identify miR-200f candidate genes were bench validation of down-regulation of the genes by *miR200-b*, associations with human disease, and significant changes in mRNA expression levels in miR200-b treated cells. While no bench validation was performed in this study, we have identified a set of miRNAs whose differential expression corresponds with deregulation of a number of significantly differentially expressed glycosyltransferases in a number of cancers (Table 7).

**Table 7.**
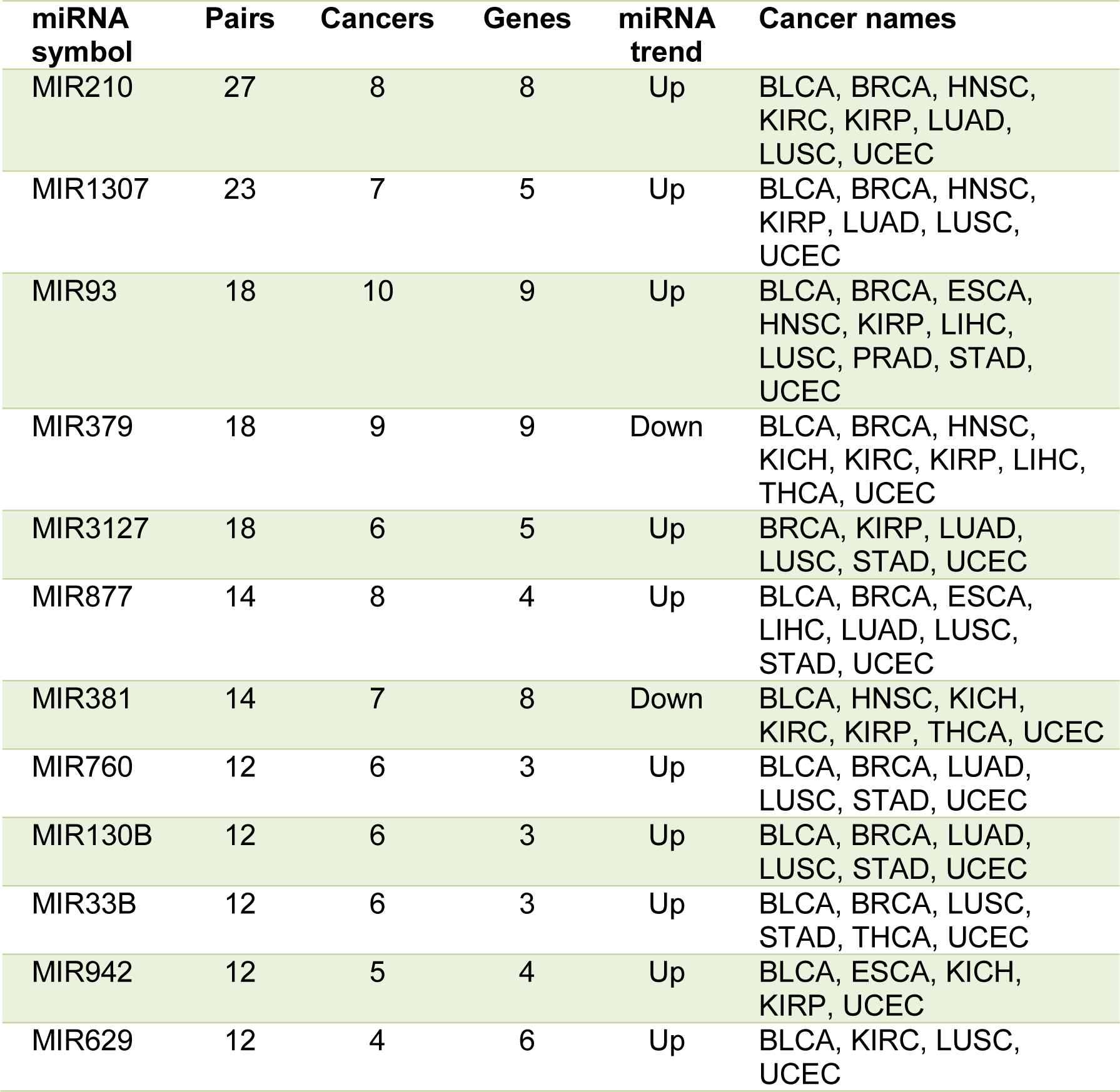
Differentially expressed miRNAs with more than 10 miRNA-GT pairs in cancer.

*miR-210* is well documented as a hypoxia regulated miRNA over-expressed in many cancers (Qin et al. 2014; Greville et al. 2016; Bar et al. 2017). *miR-210* has been found to target thrombospondin type I domain containing 7A (*THSD7A*), an important N-glycoprotein in preeclampsia (Luo et al. 2016), and hypoxic microenvironments are thoroughly categorized to change the expression of glycosyltransferases (Koike et al. 2004; Croci et al. 2014; Nonaka et al. 2014). There are two reports of *DDOST* as a predicted downregulated target of *miR-210* in lung adenocarcinoma A549 cells (Puissegur et al. 2011) and in activated *miR-210* mimic and *miR-210* TG B cells (Mok et al. 2013), but there was no other literature found linking *miR-210* regulation of glycosyltransferases in cancer. Moreover, *miR-210* is not included in the set of miRNAs identified by Agrawal et al. as belonging to miRNA/glycogene regulatory networks (Agrawal et al. 2014). In light of the prominent role of *miR-210* in cancer and the mounting evidence for miRNA and glycogene co-regulation in cancer, the co-occurrence of *miR-210* over-expression and under-expression of many GTs in several cancer types should be studied further for potential validation of *miR-210* as a regulator of GTs in cancer.

The remaining miRNAs with abundant co-differentially expressed GTs across cancers have varying levels of evidence in literature regarding roles in cancer. *miR-1307* contributes to breast cancer development(Han et al. 2019) and has been found to be upregulated in chemoresistant epithelial ovarian cancer (Zhou et al. 2015). *miR-93* promotes tumor angiogenesis through *LATS2* suppression (Fang et al. 2012), and has been reported to target *TGFβR2* in nasopharyngeal carcinoma (Lyu et al. 2014). *miR-379* expression has been found significantly decreased in breast cancer (Khan et al. 2013). *miR-3127* is a tumor suppressor in non-small-cell lung cancer (NSCLC) (Sun et al. 2014), but has been observed with oncogene activity in other cancers (Tello-Ruiz et al. 2020). *miR-877* has been upregulated in hepatocellular carcinoma (HCC) cell lines (Cai et al. 2017) and may mediate paclitaxel sensitivity through targeting of *FOXM1* (Huang et al. 2015b). *miR-381* is a tumor suppressor in colorectal cancer (He et al. 2016) and metastatic inhibitor in gastric cancer (Cao et al. 2017). *miR-760* has been found to be downregulated in many cancers (Manvati et al. 2020), including chemoresistant breast cancers (Monchusi and Kaur 2020). *miR-130B* is a prognostic indicator in pancreatic cancer (Zhao et al. 2013), and has been suggested to have an oncogenic role in prostate cancer (Fort et al. 2018). *miR-33B* inhibits stemness in breast cancer cells (Lin et al. 2015) and suppresses proliferation of HCC cells(Tian et al. 2016). *miR-942* has been significantly upregulated in HCC (Zhang et al. 2019) and induces metastasis in NSCLC (Yang et al. 2019). *miR-629* promotes pancreatic cancer (Yan et al. 2017) and is an oncogene in several other cancers, but not laryngeal cancer (Yan et al. 2017).

While the participation of these and other miRNAs in cancer is well-supported in literature, specific references to regulating GTs were not found. However, the overall evidence supporting the link between miRNAs and GTs has exploded in the last five years, resulting in the knowledge of over 80 known glycogene targets of miRNAs (Thu and Mahal 2020), much of which has been based on prediction algorithms and databases like miRanda (Enright et al. 2003) and TargetScan (Agarwal et al. 2015). Thus, the miRNA-GT targets reported here, based on prediction by miRTarbase (Huang et al. 2020), miRecords (Xiao et al. 2009), miRWalk (Sticht et al. 2018), miRTex (Li et al. 2015), DIANA-Tarbase (v6) (Karagkouni et al. 2018), and CancerMiner (Jacobsen et al. 2013), for miRNAs with co-occurring DEGTs in cancer, plus the accumulating evidence for wide-scale miRNA regulation of the glycome through glycoenzymes, provides a rationale for prioritizing these genes for validation as miRNA targets in cancer.

### Functional impact on N-glycan synthesis due to DEGTs in cancer

N-glycans share a common pentasaccharide GlcNAc2mannose(Man)3 core that can be further modified by galactosylation, GlcNAcylation, sialylation, and fucosylation (Varki 2017; Reily et al. 2019), each modification accomplished by a different enzyme. This model increases complexity and diversity, but also increases the number of potential targets by which deregulation can occur. As seen in Table 5, the number of potential glycans impacted by deregulated GTs in cancer is substantial, although the numbers reported are likely inflated by selection of an arbitrary log2FC threshold. In reality, the expression at a given time is a complex interplay of numerous environmental stimuli and regulatory effectors, both attributable to sample biology and effect from condition (cancer). The analysis method used can be confidently interpreted such that a reported fold change associated with a given *P* value after adjustment for multiple testing is a true non-zero change for the selected FDR, but it cannot dictate how meaningful a change of a given effect size may be. However, the log2FC thresholds or rankings based on such are imposed to identify some subset with the largest effect size and therefore more likely to be meaningful. Figure 7 shows the relationships between cancer type, direction of change, gene, and residue impacted in implicated glycans for a log2FC magnitude threshold of 3.0. In the 12 different cancer types shown, more DEGTs involved in N-linked glycosylation have an expression change of “Up” or increase than “Down”. The nine DEGTs identified at |log2FC| > 3.0 represent multiple families. Residues impacted are largely affected by a single enzyme in cancer, but may be compensated for by other enzymes not differentially expressed, allowing the glycan profile to resist changes due to aberrant glycosylation. The fucosyltransferases, however, while having high redundancy for residues, are also substantially impacted in cancer, and could therefore have a multiplied effect on fucosylation in cancer. The separation of residue impact by enzyme or enzyme family can be easily seen, where most residues are affected by a single differentially expressed enzyme, but the fucosyltransferases are each impacting three potential residues, three of which are acted upon by *FUT3*, *FUT6*, and *FUT9*. This is consistent with reports of increased fucosylation in multiple cancer types (Keeley et al. 2019), lending merit to the approach used to identify DEGTs of biological significance. Therefore, it is suggested that these enzymes, as well as the remaining five N-glycosyltransferases reported here, be prioritized for study as possible cancer biomarkers.

### DEGTs as known cancer biomarkers

EDRN overlap analysis revealed that all GTs in the EDRN biomarker dataset were indicated in prostate cancer. However, literature review found that several of these biomarkers were also implicated in various other cancers. For example, *CSGALNACT1* is implicated mostly in prostate cancer, but also serves as a prognostic biomarker in multiple myeloma (Qi et al. 2020). *GALNT7* is favorable as a prognostic biomarker in colorectal cancer and gliomas, but has unfavorable results in pancreatic cancer with a lower survival rate [https://www.proteinatlas.org] (Uhlen et al. 2015; Hua et al. 2018). *STT3A* has also been recognized as a diagnostic panel biomarker for thyroid cancer, and *ALG10* autoantibodies have been implicated as potential diagnostic biomarkers for breast cancer, consistent with the data in EDRN (Zhong et al. 2008). In addition to being a known prostate cancer biomarker, *GALNT3* is a favorable prognostic biomarker in cervical cancer and unfavorable in head and neck cancers [https://www.proteinatlas.org/] (Uhlen et al. 2015). Finally, *OGT* has also been implicated in prostate cancer and has been reported to be a poor prognostic biomarker in colon cancer (Wu et al. 2020). In the FDA approved biomarker datasets *EXT1* in breast cancer was the only biomarker which overlapped with the DEGT dataset.

As indicated above, several DEGTs have been reported as effective biomarkers in literature; however, a comprehensive knowledgebase to curate and store a list of DEGTs extracted from publications is needed for further evaluation. Five DEGTs were randomly selected and were found to be reported as biomarkers ranging from prognostic biomarkers to risk assessment biomarkers. *LARGE1* is a prognostic biomarker in epithelium derived cancers (Liu et al. 2021). *PYGB*, indicated as a potential high-value GT in several experiments described herein is a prognostic biomarker in ovarian cancer (Zhou et al. 2019). HAS family members (*HAS1*, *HAS2*,) have been well documented as prognostic and diagnostic bladder cancer biomarkers (Kramer et al. 2011). *B3GALT4* has been implicated as a prognostic biomarker in colorectal cancer(Zhang et al. 2018a), and *EOGT* is an unfavorable prognostic biomarker in renal cancer [https://www.proteinatlas.org] (Uhlen et al. 2015). A planned extension of this foundational DEGT dataset is to build a biomarker resource using the BEST (Biomarkers, EndpointS and other Tools) guidelines (2016). The resulting DEGT dataset will be hosted at https://data.oncomx.org/cancerbiomarkers under a tab called “DEGT biomarker data.”

### Integrating findings toward a high-value list of potential glycosyltransferase cancer biomarkers

The high-value glycosyltransferases identified from the various analyses described above represent a set of genes with at least one (and frequently many) experimental evidences of a cancer association. In many cases, reports of some aspect of these genes in cancer are described in the literature, suggesting that genes identified by these methods have the potential to be biologically meaningful with respect to their presence in cancer. Table 8 summarizes the high-value candidates reported throughout the study. Unsurprising based on the number of mentions throughout this work, *PYGM* has the most appearances across evidence types with six, with all but glycan impact and biomarker status. *MFNG*, *UGT2B7*, *FUT2*, *FUT6*, and *GALNT3* have three evidences each, and the remaining GTs in the table below have two evidences each, except *GALNT5* with a single evidence of disease literature in the high-value list. Of note, *GALNT5* is SDE and is predicted to be targeted by SDE miRNAs in multiple cancers, but is excluded from the high-value list in these categories because of a lower rank. In future implementations, iteration of all analyses performed herein using complete lists of high-value genes (as opposed to just the top ranked) could highlight interesting links between data types and approximate a better overall picture of the role of GTs in cancer.

**Table 8.**
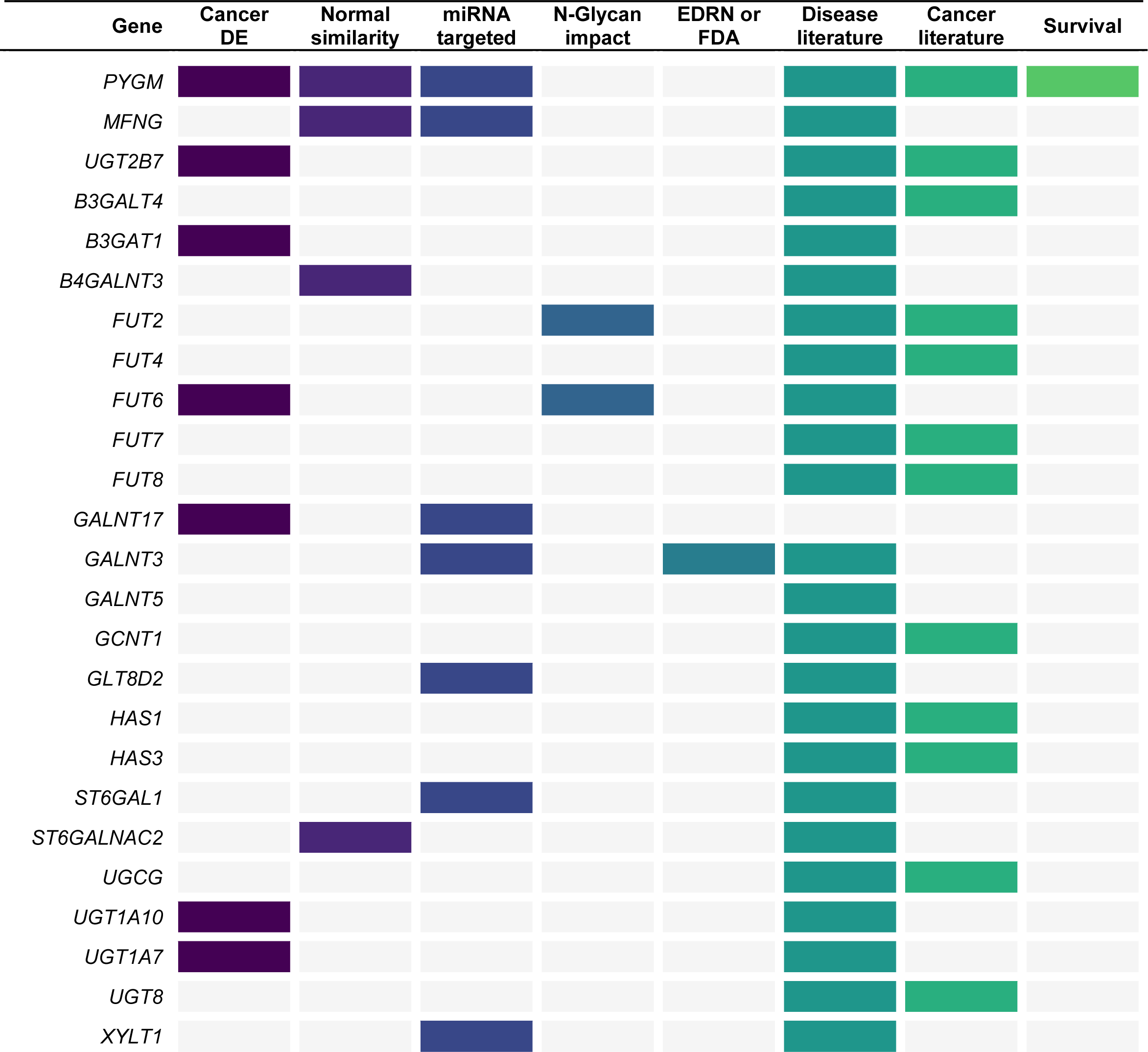
Accumulation of evidence for high-value glycosyltransferases in cancer.

### Analysis of survival in cancer for high-value DEGTs

Of the 39 gene-cancer pairs evaluated for survival based on relative gene expression in tumor samples, five showed significant differences in overall survival between high and low expression groups (Figure 8). High expression of two genes from the N-acetylgalactosaminyl transferases (GALNTs) family was significantly associated with worse survival probability. *GALNT15* and *GALNT16* produce the proteins N-acetylgalactosaminyltransferase 15 and 16, respectively, and serve an enzymatic function in the transfer of N-acetylgalactosamine (GalNAc) from UDP-GalNAc to the serine and threonine residues on O-linked glycoproteins (Peng et al. 2010). *GALNT16* is grouped in subfamily Ib of the GALNTs and is nearly identical in sequence to other GALNTs for subfamily Ib in the catalytic and lectin domains, while *GALNT15* does not group into any subfamily based on phylogenetic analysis and intron positions (Bennett et al. 2012).

Although our findings indicate a relationship between *GALNT15* expression and prognosis in endometrial cancer, *GALNT15* has not been extensively studied in endometrial cancer. The genetic variant rs2102302 in *GALNT15* was found in a study of Spanish patients with colorectal cancer, although the variant was not validated in a second, larger group (Abuli et al. 2011). Further evidence is needed to elucidate how *GALNT15* relates to patient prognosis in endometrial cancer and cancer overall.

Similarly, *GALNT16* has not been extensively studied in bladder cancer; however, our findings of poorer survival for increased expression of *GALNT16* in tumor tissue from the BLCA study group is consistent with findings for higher expression of *GALNT16* in other cancers. *GALNT16* has been implicated in pancreatic cancer and may be downstream of pancreatic cancer progression through *GAL1*, showing increased *GALNT16* mRNA expression in recombinant Gal1 treated human pancreatic tumoral cells (Orozco et al. 2018). Certain polymorphisms in *GALNT16* have been associated with increased breast cancer risk when evaluated in Han Chinese breast cancer patients (Wu et al. 2019). While higher expression of *GALNT16* has been found in breast cancer tissue compared to normal, and higher expression has been related to increased risk through Kaplan-Meier analysis (Wu et al. 2019), the DE analysis conducted for this study did not find *GALNT16* to be significantly differentially expressed in breast cancer. This could suggest that possible association of *GALNT16* with cancer is population specific, or that the analysis method used in the DE study lacked the sensitivity to identify such a relationship. However, previous survival analysis in patients from the TCGA BLCA study confirm our analysis of decreased overall survival with higher *GALNT16* mRNA expression [http://www.proteinatlas.org/] (Uhlen et al. 2015).

Interestingly, from the differential expression analysis, these GALNTs were typically found to be under-expressed in cancer when comparing adjacent normal tissue, with the exception of *GALNT16* in KICH. Note that survival analysis is only comparing expression among tumor samples, and therefore is showing that higher expression among tumor samples is associated with worse prognosis. Although not able to be deduced from the experiments conducted for this study, one possible explanation is that individuals who have high expression of *GALNT15* or *GALNT16* overall may have worse prognosis, even though these GTs are expressed higher in their adjacent normal tissue than in the tumors themselves. Further examination is required to determine the possible relationship between these findings.

The glycogen phosphorylase genes *PYGM* and *PYGB* were both found to be significantly linked with survival, with higher expression of *PYGM* linked with worse survival in the UCEC study group and higher expression of PYGB linked with worse survival in the BLCA study group. Both the muscle form, encoded by *PYGM*, and the brain form, encoded by *PYGB*, catalyze glycogen to glucose-1-phosphate in carbohydrate metabolism (Gelinas et al. 1989; Llavero et al. 2019). *PYGM* has been studied extensively for its role in McArdle disease, however, little is known about how expression of *PYGM* relates to cancer prognosis. Recently, *PYGM* was evaluated as a biomarker of head and neck squamous cell carcinoma, with high expression of *PYGM* linked with worse survival (Jin and Yang 2019). *PYGB* has been implicated in cancer cell metabolism of glycogen for increased survival, as well as for poorer prognosis when highly expressed in specific cancer types. In glucose-resistant pancreatic cancer cells, downregulation of *PYGB* led to increased cell death upon glycogen starvation(Philips et al. 2014). *PYGB* phosphorylation increased with higher expression of KIAA1199, a marker of poorer prognosis in gastric cancer (Terashima et al. 2014; Jia et al. 2017). Finally, increased expression of *PYGB* was identified and linked with poorer prognosis in both ovarian cancer and hepatocellular carcinoma(Zhou et al. 2019; Cui et al. 2020). Previous survival analysis of *PYGB* in the BLCA study group showed a similar trend of higher mRNA expression linked with worse survival (Uhlen et al. 2015).

Similar to the GALNTs, *PYGM* and *PYGB* were identified by DE analysis to be under-expressed in most cancers compared to adjacent normal samples, but inked with worse survival in the high expression groups. However, *PYGB* was reported to be over-expressed in cancer with worse survival in the high expression group. While it is important to remember that DE analysis and survival are looking at two different comparisons, tumor vs. normal and tumor high expression vs. tumor low expression, respectively, this observation could suggest that increased expression of *PYGB* in hepatocellular carcinoma is both diagnostic and prognostic. Further study is necessary to determine a mechanistic relationship between these two observations.

*POMGNT2*, the final GT identified with expression groups to be significantly different in overall survival for the KIRC study group, encodes protein O-linked-mannose beta-1,4-N-acetylglucosaminyltransferase 2, one of multiple glycosyltransferases that catalyze the glycosylation of α-dystroglycan (Manzini et al. 2012). High expression of *POMGNT2* has been identified in brain, muscle, and kidneys, and mutations can cause a range of presentations from mild muscular dystrophy to Walker-Warburg syndrome (Manzini et al. 2012; Endo et al. 2015). In cancer, laminin binding to cell surface proteins, including dystroglycan, increases tumor growth, and disruption of α-dystroglycan has previously been linked with increased mortality in renal clear cell carcinoma (Akhavan et al. 2012; Miller et al. 2015). Downregulation of other GTs responsible for glycosylation of α-dystroglycan has also been linked with increased mortality in patients with renal clear cell carcinoma(Miller et al. 2015). The findings presented herein of low expression of *POMGNT2* linked with worse prognosis align with previous survival analysis of *POMGNT2* expression in the KIRC study group (Uhlen et al. 2015). Moreover, analysis of *POMGNT2* differential expression identified significant under-expression, suggesting that low expression could be both a diagnostic and prognostic indicator.

### Limitations of selected analysis methods and impact on reported results

The rationale for the differential expression method employed in this analysis was based on the adoption of the same or similar methods described throughout biomedical literature during the time of work performed, through to the time of publication. Specifically, DESeq2 has been commonly used to compute differential expression in disease conditions compared to some reference condition, including ulcerative colitis (Lee et al. 2020), aortic aneurism (Zalewski et al. 2020a), lung adenocarcinoma (Xiong et al. 2020), COVID-19 (Ramesh et al. 2021), and more (Applebaum et al. 2020; Nali et al. 2020; Zalewski et al. 2020b). Common criteria used to call differential expression includes *P* < 0.05 (after adjustment for multiple testing) and a log2FC magnitude of at least 1.0 (Andrade et al. 2020; Colpo et al. 2020), although the exact parametric space used across studies is variable with respect to *P*, fold change, and FDR levels (Applebaum et al. 2020; Grabski et al. 2020; Zalewski et al. 2020b). One major limitation of the study described herein was in using the default parametric space for the DESeq2 implementation, which includes a less stringent FDR α = 0.1, expected to result in a higher degree of false positives than desirable. When testing tens of thousands of genes and identifying thousands to be differentially expressed, this may result in erroneous identification of up to hundreds of genes. For this reason, very restricted subsets of those DEGTs having the highest relative estimated fold changes were advanced to designation as high-value candidates, but the threshold used was mostly arbitrary, selected primarily based on the number of candidates resulting from the threshold, not the quality of high-value DEGT candidates identified. In this case, because of the small proportion of selected high-value DEGTs compared to the set of all DEGTs, loss of significant results is more likely than including falsely identified significant results. Identifying subsets based on the distribution of results instead of selection of an arbitrarily discrete count would likely improve the overall robustness of high-value identification.

Another limitation involves the use of paired data (coming from the same sample) without controlling for possible patient-related-similarities between these samples (as compared to samples coming from other patients). At the onset of analysis, the software packages investigated were not configured to deal with paired data; however, options now exist within DESeq2 and other packages to do exactly this. Updating α and specifying pairing in the model would greatly improve the confidence of the resulting set of DEGs reported. Modeling of clinical features may also improve the ability to accurately identify DEGs, as well as derive findings of biological and clinical relevance.

In addition to optimizing the parametric space and modeling assumptions for better results, tests could be employed to analyze the performance of the pipeline on a variety of gene sets to better inform the interpretation of differential expression of the gene set of interest (GTs). Robustness testing could be employed to investigate the quality of the results from independent datasets (Sim et al. 2017), for example, from a cancer dataset not provided by TCGA but of the same type. Permutation testing could be employed across randomly sampled subsets of data to define DEGs across permutations. DE analysis could also be analyzed with different software to define DEGs only as those in the intersection of results from different strategies. Moreover, new methods are constantly being developed to address issues related to the quality of differential expression analysis and improve the identification of DEGs. Recent methods have been proposed to overcome the issue of threshold decisions (Sim et al. 2017), and to employ the knowledge of a gene’s prior probability of DE to better predict that gene’s differential expression across diverse biological experiments (Crow et al. 2019).

Enrichment analysis was performed using a cumulative binomial test based on the methods described by Mi and Thomas (Mi and Thomas 2009), previously used in PANTHER gene enrichment analysis. However, review of current methods revealed a change in the default method used by PANTHER to the Fisher’s Exact test, corresponding to the methods used by a number of GO enrichment tools (Rivals et al. 2007). Enrichment was not re-analyzed for all GT and family comparisons, but Fisher’s Exact *P* values were calculated as validation for the subset of GT families identified to be over-expressed in various cancers.

### Data provenance capture and FAIRness

The implementation of BCOs in the GlyGen data integration and standardization workflow was critical to disambiguating potential errors inherent to the manually generated input dataset and other errors introduced in post-processing steps, preventing diminished value of the database due to incomplete provenance description (Wooley et al. 2005). With subsequent partial automation and updates to the pipeline and resulting dataset, provenance details can now be supplied for the entire pipeline, including both preliminary processing (the part described in this study) and steps taken by GlyGen to generate and host the final product file. Versioning of the object associated with this dataset and all updates thereafter will automatically occur every time the object is modified in the database, and pertinent details will be captured in the corresponding BCO in JSON format. The record of the executed processes and the defined parametric space together, accompanying the product of the processing, is expected to drastically improve the interoperability and reusability components of the FAIR Data Principles (Wilkinson et al. 2016), as well as overall reproducibility.

### Outlook and future directions

The strategy described herein presents two pipelines: one for the definition of the list of human glycosyltransferase enzymes, and the second for the analysis of differentially expressed genes in cancer, starting from RNA-Seq derived counts data and comparing across data from multiple types of experimental expression data. Many datasets generated for this study can be used to quantify expression for other genes or diseases, and are expected to be leveraged in the near term to study other glycoenzymes toward a more comprehensive understanding of glycosylation in cancer. Differential expression pipeline results for all cancers, and many of the other experimental datasets described throughout, are integrated as part of the OncoMX database for cancer biomarkers and related genes. The subset of identified high-value target GTs can be used in additional characterization studies like the ones described above, and updates to the expression calling portion of the analysis could generate more robust lists of high-value biomarker targets.

### Conclusion

Glycosylation is increasingly implicated in cancer development and progression (Borzym-Kluczyk et al. 2012; Munkley and Elliott 2016; Burchell et al. 2018; Reily et al. 2019), but the underlying mechanisms driving this relationship necessitate substantial additional exploration (Thomas et al. 2020). Specifically, glycosyltransferases have been identified in cancer in various roles and scopes: individual GTs in individual cancer types (Miller et al. 2015), individual GTs in multiple cancer types, groups of GTs in individual cancers (Carlini et al. 2005; Barthel et al. 2008; Gupta et al. 2020), and groups of GTs in multiple cancers (Petretti et al. 2000; Ashkani and Naidoo 2016). Critical to the understanding of the role these enzymes may play in cancer is the adequate and accurate definition of the class of GTs and the enzymes belonging to that class. Recent efforts to better define and characterize the normal functions of GTs (Hansen et al. 2015; Joshi et al. 2018), including efforts described in this work, greatly improve the potential for meaningful interpretation of their impact in diseases, including cancer.

A survey of differential expression of GTs in cancer identified that DEGTs are abundant in kidney, breast, and lung cancers but are present in most cancer types, and that most GTs identified with higher levels of change are differentially expressed in many cancers, albeit at different levels and often in different directions. Limitations of the analysis method used include a less-stringent FDR and pair-blind analysis, likely resulting in a higher number of reported DEGs with lower *P* values, even after correction by multiple testing. However, literature review of high-value DEGTs defined by ranking change suggests the analysis provided herein was capable of identifying candidate genes with real biological value. Future pipeline improvements to accommodate identification at a lower FDR and leveraging newer software developments for substantially improved treatment of paired data are expected to greatly improve the quality of hypotheses generated by this approach.

While significant differential expression provides one means of interrogating the importance of GTs in cancer, technological advances combined with global initiatives and a general philosophical consensus toward data sharing enables integration of multifaceted information types with the lowest barriers yet encountered in genomics. Although many additional data types are available, the analysis discussed here leveraged various modes of expression data, including normal (assumed healthy) expression across human and mouse, miRNA-Seq-derived differential expression, and automatically mined literature reporting differential expression in cancer. Nine DEGT/cancer pairs, each gene of which has literature of relevance to cancer, were identified to be high-value based on comparison of normal expression across organisms and differential expression in cancer. Numerous differentially expressed miRNAs targeting DEGTs in cancer were identified, an observation with increasing coverage in the scientific literature. Automated literature coverage was relatively low, partially reflecting the rapid increase of available literature on the topic of GT expression in cancer during the time this study was performed, but also suggesting room for improvement to the text extraction algorithms employed. Overlapping the GlycoTree annotation map with the high-value DEGTs, a set of residues and N-glycans were identified based on the expectation of impacted synthesis due to differential expression of at least one of its enzymes. Overlap of GTs with existing curated biomarker sets resulted only in partial maps, with a small number of GTs identified as biomarkers but for different cancer types than those reported in the high-value subset of this analysis. Finally, high-value DEGTs identified from across all experiments were ranked, with the highest ranked GT/cancer pairs subjected to survival analysis. Five GTs from this set were found to have significantly different survival outcomes between high and low tumor expression groups. Based on the aggregated evidence across experimental data, along with literature support for cancer involvement, the high-value DEGTs are presented as candidate genes to be further studied for roles as cancer biomarkers.

The process of identifying cancer biomarkers is immensely costly, both in terms of time and monetary resources. Combined analysis of multiple diverse information types can assist in the identification of high-value candidate biomarkers, prioritizing candidates for downstream study and validation. However, significant barriers to data access and interoperability are rampant across the domain of omics-level information. The OncoMX cancer database and knowledgebase for cancer biomarker and related evidence was designed to address these issues and enhance the potential downstream utility of the results of the analysis presented herein. Although the scope of OncoMX is vast and extends far past the glycosyltransferase application described in this work, various OncoMX features and datasets were designed based on this use-case. Data generated for and by this work is available from the OncoMX data portal. It is expected that an easily accessible, unified, highly integrated cancer database including biomarker data and various evidence types can promote reusability of data and improve the process by which biomarker hypotheses are generated, as demonstrated by this study.

## Methods

### Data retrieval and curation

Proteomes and gene and protein lists were first retrieved from UniProtKB/Swiss-Prot (UniProt 2019) (UniProt release 2016_01) accessed February of 2016, and updated October of 2020 (UniProt release 2020_04). The full set of proteins comprising the human proteome was retrieved through API using the following query: organism:9606+AND+reviewed:yes+AND+proteome:UP00000564. The subset of human proteins annotated with the “Glycosyltransferase” UniProt keyword were retrieved with the query: “organism:9606+AND+reviewed:yes+AND+keyword:“Glycosyltransferase [KW-0328]”+AND+proteome:UP00000564,” and those specifically annotated by the Carbohydrate Active Enzymes database (Lombard et al. 2014) (CAZy, http://www.cazy.org/) were retrieved with the query: “organism:9606+reviewed:yes+proteome:UP000005640+database:(type:cazy glycosyltransferase).” GTs annotated by the Consortium for Functional Glycomics (CFG) functional glycomics gateway (http://www.functionalglycomics.org/) were scraped from the web results for the “Query by Multiple Criteria” for any classification and linkage for human enzymes. A slim was created using QuickGO [https://www.ebi.ac.uk/QuickGO/] to retrieve proteins annotated with molecular function GO term, “GO:0016757 transferase activity, transferring glycosyl groups,” and relevant child terms. Terms not involved in glycan synthesis or attachment were excluded. The resulting list of annotated GTs was filtered to remove any duplicate references resulting from cross-database redundancy or secondary accessions. InterPro family (Mitchell et al. 2015) annotations were retrieved using EBI API with query term “glycosyltransferase.” Pfam domain (Finn et al. 2016) annotations were then retrieved for all genes in the list, and manually curated to produce a complete list of glycosyltransferase-related annotations. Figure 2 is a schematic representation of the GT list generation pipeline. Supplemental Table S1 displays the full list of automatically captured genes (including hits later excluded) and corresponding annotations retrieved through the above-described workflow. Supplemental Table S2 displays the final enzyme list following manual curation of entries. Additional functional annotations from the lists of GTs published by Hansen et al. (Hansen et al. 2015) and Joshi et al. (Joshi et al. 2018), including GT fold type, glycosylation pathway, synthesis step, donor substrate, isoenzyme and regulation status, cellular localization, and protein type, have been appended to all corresponding GT entries (Supplemental Table S3).

Pre-processed read counts and FPKM gene quantification for all cancers were retrieved from the NCI Genomic Data Commons (GDC) data portal (Grossman et al. 2016) for TCGA studies [https://www.cancer.gov/tcga] with samples of both sample type codes 01 Primary Solid Tumor and corresponding 11 Solid Tissue Normal. FPKM normalized read count values were downloaded only for cancer samples, along with corresponding metadata file and clinical information file downloaded for each cancer. Read counts were analyzed as described by the original BioXpress pipeline (Wan et al. 2015) (February 2016) and updates (Dingerdissen et al. 2018), accessed from the OncoMX (Dingerdissen et al. 2020) data portal at https://data.oncomx.org (April 2020) to determine differential expression of the identified glycosyltransferase genes between matched tumor and adjacent normal pairs for each cancer.

The set of human and mouse pairwise orthologs were retrieved from the Orthologous MAtrix (OMA) database (Altenhoff et al. 2018) REST API (Kaleb et al. 2019) for the set of all genes with a 1:1 relationship, such that both the human and mouse genes in a given pair were each other’s mutually exclusive ortholog for the two species. A list of mouse GTs was retrieved from GlyGen (York et al. 2020) data portal at https://w3id.org/biocompute/portal/GLY_000030. Expression calls were retrieved for assumed healthy human and mouse individuals for the set of mammalian anatomical structures with data for both from Bgee (Bastian et al. 2020) through OncoMX (Dingerdissen et al. 2020) in April 2020.

miRNA target information was extracted from supplemental information in Hu et al. (Hu et al. 2018) for a subset of miRNAs significantly differentially expressed in multiple cancers with target genes identified to be significantly differentially expressed in the same cancer type. Differential expression of these miRNAs was retrieved from OncoMX at https://w3id.org/biocompute/portal/OMX_000022.

A file of automatically mined text relationships for GT expression in disease generated by custom implementation of DEXTER (Gupta et al. 2018) software for Disease-Expression Relation Extraction from Text was provided directly by collaborators. A second custom prepared file of automatically mined text relationships for expression of all genes in cancer was generated for and retrieved from OncoMX athttps://w3id.org/biocompute/portal/OMX_000004.

Existing cancer biomarker details were retrieved from the lists of public Early Detection Research Network (EDRN) biomarkers and FDA approved cancer biomarkers for breast, colon, lung, ovary, prostate, and melanoma, available at https://w3id.org/biocompute/portal/OMX_000019, https://w3id.org/biocompute/portal/OMX_000003, https://w3id.org/biocompute/portal/OMX_000026, https://w3id.org/biocompute/portal/OMX_000027, https://w3id.org/biocompute/portal/OMX_000028, https://w3id.org/biocompute/portal/OMX_000029, and https://w3id.org/biocompute/portal/OMX_000030, respectively, in March 2021.

Annotated glycans from GlycoTree (Takahasi 2003) were retrieved from the GlyGen (York et al. 2020) data portal from object GLY_000284 in April 2020. An updated version of the same file currently in the GlyGen test server was provided directly by collaborators in September 2020.

### Differential expression of glycosyltransferases

Differential expression was pre-computed for all genes as defined in the BioXpress pipeline, previously described by Wan et al. (Wan et al. 2015) and Dingerdissen et al. (Dingerdissen et al. 2018) In brief, read count data were retrieved for patients with RNA-Seq derived counts for tumor and solid normal samples as identified using the advanced search queries “files.analysis.workflow_type in [“HTSeq - Counts”] and files.data_type in [“Gene Expression Quantification”] and cases.samples.sample_type in [“Primary Tumor”] and cases.project.program.name in [“TCGA”]” and “files.analysis.workflow_type in [“HTSeq -Counts”] and files.data_type in [“Gene Expression Quantification”] and cases.samples.sample_type in [“Solid Tissue Normal”] and cases.project.program.name in [“TCGA”]” for tumor and paired solid tissue normal samples, respectively. Ensembl IDs were mapped in counts data to gene symbols for samples from studies with more than 10 sample pairs. For each remaining study, the Bioconductor (Huber et al. 2015) DESeq2 (Love et al. 2014) package for differential analysis of count data was used to analyze the gene counts data from TCGA (generated with python package HTSeq (Anders et al. 2015) in GDC pipeline (GDC 2020)) using tumor/normal status as categories for defining the differential analysis. Significance was reported by Wald test *P* value, adjusted for multiple testing by the Benjamini-Hochberg procedure (Benjamini and Hochberg 1995). Results were ordered by adjusted *P* value and filtered for rows containing log2FC values of “NA.” Individual sample log2FC were calculated and results were loaded to the final table, mapped to corresponding gene, protein, disease, and tissue annotations.

The complete file with differential expression calculations for all genes in cancer was filtered to keep only those genes corresponding to UniProtKB/Swiss-Prot reviewed entries. The subset of genes with adjusted *P* < 0.05 for differential expression in a given cancer defined a working set of significant genes for each cancer type. Significant genes were further filtered for each glycosyltransferase in the curated list to define a set of significant GTs for each cancer, and aggregated to define the set of all GTs significantly differentially expressed in at least one cancer and across all available cancer types.

Resulting tables were transformed for compatibility with visual tools. DEGTs were filtered for increasing log2FC thresholds (in magnitude of 1.0, 2.0, 2.5, 3.0, 3.5, and 4.0) to enable examination of various cutoffs.

For each log2FC threshold cutoff, summary counts were compiled for (1) each GT with respect to the number of cancers in which that gene was increased, decreased, or had no reportable change, and (2) each GT with respect to the number of patients for which that gene was increased or decreased across all cancers. All identified DEGTs with adjusted *P* < 0.05 and log2FC magnitude of at least 1.0 can be found in Supplemental Table S4. Supplemental Table S5 summarizes DEGTs by the total number of cancers in which each GT was reported to be differentially expressed, as well as the subset counts for cancers in which each GT was over- or under-expressed.

### Enrichment analysis and statistics

Enrichment significance of gene and family counts was determined using a cumulative binomial test based on previously described and implemented methodology (Mi and Thomas 2009; Dingerdissen et al. 2013). In short, the ratio of the number of genes in the set of all GTs to the number of all genes in the human genome defined the expected ratio of the number of significantly differentially expressed GTs in a given cancer to the number of all significantly differentially expressed genes in that cancer (significant differential expression defined by BH adjusted *P* < 0.05 and |log2FC| ≥ 1.0). The sample frequency was compared to the background frequency such that probabilities for under-representation of DEGTs in a given cancer were equivalent to the cumulative probability of detecting any count less than or equal to the observed count based on the expected count.

Similarly, probabilities for over-represented significantly differentially expressed GTs in some cancer were equivalent to the cumulative probability of detecting any count greater than or equal to the observed count based on the expected count. Enrichment for differential expression of families of GTs in individual cancers was calculated in the same way, replacing the set of all GTs with the subset of all GTs in a given family.

Enrichment of differentially expressed GTs in cancer against the background of all differentially expressed human genes in all cancers was calculated using the above described methodology with the resulting *P* value adjusted for multiple testing by the Bonferroni correction such that a significant finding would have *P* < 0.05/(number of cancers). Enrichment of individual GT families by differential expression in all cancers against the background of all genes in all cancers was similarly calculated with each family *P* value adjusted by Bonferroni correction for the number of cancer types.

### Defining orthologs

Human-mouse orthologous GT gene pairs were defined by overlapping the subset of 1:1 pairwise GT orthologs from OMA (Altenhoff et al. 2018) with the set of mouse GTs retrieved as object GLY_000030 from GlyGen.

### Analysis of healthy expression in human and mouse

Normal expression calls and quality assessments for human and mouse were generated as described in Bgee documentation (Bgee 2020) and summarized in the description of OncoMX by Dingerdissen et al. (Dingerdissen et al. 2020) Additional processing considerations specific to the integration into and custom presentation of the data in OncoMX can be found in the data supplement corresponding to Dingerdissen et al. (2020) (Dingerdissen et al. 2020) and in the Bgee GitHub pages describing the OncoMX collaboration. Briefly, expression categories were defined by comparing the rank score of the expression call (for the subset of genes with present calls) for a given gene in a given tissue with the minimum and maximum ranks of that same gene across all tissues in which it was found to be expressed. Genes were called with “HIGH” expression in tissues where that gene has a low expression rank score compared to the range of rank scores across all tissues. Thresholds for HIGH, MEDIUM, and LOW categories were dependent on each gene/tissue distribution by considering the difference between the maximum and minimum ranks for each gene in a given tissue and dividing the difference by the number of levels (three). In cases having a difference between maximum and minimum less than or equal to 100, the level was always designated “HIGH.”

### Calculating similarity impact score

A simple score was devised to facilitate numerical summarization of the similarity of behavior of genes across multiple cancer types in a fashion amenable to visualization. For each normal expression qualification, a score of 1, 0, or -1 was added for calls of HIGH, MEDIUM, and LOW, respectively. A score of -2 or 2 was added for differential expression trends of up or down, respectively. A weight of -10 was added to the score for absent calls to promote dropout of samples when aggregating scores across multiple cancers. Supplemental Table S6 shows a color-coded matrix with all possible score combinations (excluding absent calls).

### Filtering and mapping of multifaceted data types

The list of miRNA-target associations was first modified to create a new list with individual rows for each miRNA-target pair. Unique pairs were filtered such that only miRNA-target pairs including a GT were kept.

### Identifying high-value targets

For the set of all differentially expressed GTs, high-value GTs were defined as the top 40 GT-cancer associations when ranked by magnitude of log2FC among the subset of significant DEGTs (*P* < 0.05 after BH adjustment).

For the set of GTs with healthy expression across orthologs, differential expression and normal expression tables were joined such that each row representing a gene-cancer pair was mapped to healthy expression for that gene using Uberon anatomical entity terms to relate all cancer and tissue types. ntries were filtered for the subset of rows displaying similar expression calls for human and mouse with opposing differential expression trend in cancer. Although possible that any change in expression may be meaningful, the choice to include those most different (decreased expression for HIGH-HIGH profiles and increased expression for LOW-LOW profiles) was based on the hypothesis that a significant decrease in a normally highly expressed gene or a significant increase in a normally lowly expressed gene may be both more detectable and more impactful.

Tables containing the miRNA-GT target pairs and the DEGTs analyzed from TCGA mRNA-Seq data through BioXpress were merged by UniProt accession, removing any rows for GTs without targeting miRNAs. The resulting intermediate table was mapped to the miRNA-Seq-derived differential expression results using the HGNC mapping file to map alias symbols reported from the target file to the approved HGNC miRNA symbols. Headers were renamed for consistency, and the resulting file was filtered for miRNA-GT target pairs with opposing directions of change such that if GT expression was increased, miRNA expression was decreased, and if GT expression was decreased, miRNA expression was increased. This relationship was required due to the negative regulatory nature of miRNAs silencing the expression of their targeted sequences. Resulting miRNA-GT target pairs were filtered at different log2FC thresholds. Counts for all GTs targeted by at least one miRNA were summed with respect to the total number of cancers with a reported miRNA-GT target pair, the total number of miRNAs targeting the GT across the entire dataset, and the subset of each for which the gene and miRNA had the same or different trends.

A list of potential human GTs from a pilot iteration of the GT definition process described above was provided to collaborators and developers of DEXTER (Gupta et al. 2018). An initial test set of 20 sentences retrieved from a preliminary query (including terms related to gene expression in human disease) performed on MedLine abstracts was provided back for manual review and annotation. Sentences were flagged as a true hit (keep), false hit (discard with rationale to help refine the parametric space), or maybe (signifying additional discussion). Through iteration over test sets and improvements made to the algorithm, sentences were retrieved and categorized based on the structure and the evidence contained within the sentence, and annotated by an action of expression or activation. The final form of the dataset filtered for the current set of 222 enzymes resulted in 234 literature evidences of GTs in disease. Additional details on the technical implementation and optimization of DEXTER using the GT training set are included in the DEXTER publication by Gupta et al. (Gupta et al. 2018). The gene expression literature mining workflow using DEXTER was later updated and extended in scope to include all human genes, but limited exclusively to cancers..

Overlap analysis of DEGTs and existing reported EDRN and FDA biomarkers was performed by filtering each biomarker dataset for entries with corresponding GT gene symbol and identifying conserved gene-disease pairs across the DEGT and biomarker datasets. PMIDs were manually reviewed to verify the indicated organ reported by the biomarker file and interrogate whether other organs were under consideration.

### Glycan impact

The table of glycans with annotated enzymes and residues was filtered for entries corresponding to the set of 222 human enzymes. The resulting table was merged with differential expression results such that each DE line was appended with the corresponding glycan annotations. Headers were changed for consistency and duplicate entries were removed. Merged DEGT-glycan annotation dataset was filtered for variable log2FC thresholds. Counts were summarized with respect to gene, glycan, and cancer.

### Survival analysis

FPKM values were converted to TPM values using the formula derived previously (Pachter 2011):

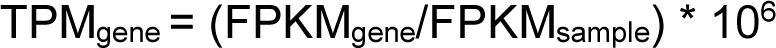

Each sample was mapped to patient ID and corresponding clinical information using the clinical and metadata files provided by the GDC data portal with customized python scripts available at https://github.com/hmdinger/glycosyltransferases-in-cancer. Gene symbols were mapped to UniProtKB/Swiss-Prot reviewed entries and unmapped genes were removed. Mapped gene expression data was then manually inspected and formatted for survival analysis. A column was created that indicated death as 1 and no death as 0. A second column was created that listed the number of days survived if death occurred or the number of days until final contact with the study if no death occurred. Median TPM value was used to differentiate high and low expression groups to account for very lowly or very highly expressed outliers, and a new column assigned each sample to either a high expression group or a low expression group. Samples were only included that contained a time interval > 0 and did not have a TPM value equal to the median TPM value. If more than half of the samples for a given TCGA study reported TPM values of zero, samples were only included in survival analysis if the sample’s TPM value > 0 for that gene.

All survival analysis statistics and Kaplan-Meier graphs were generated with R 4.0.2 (Team 2020). Specifically, Kaplan-Meier curves were generated with the survival library (Therneau 2020), and log-rank tests to generate *P* values were generated with the survminer library (Kassambara 2020).

### Visualizations

Overall workflow and computational pipeline diagrams were created in Lucidchart (Lucidchart 2020). Heatmaps and barcharts were produced using R(Team 2020) libraries ggplot2 (Wickham 2016), dplyr, viridis, and tidyr. Anatomical diagrams were produced using R(Team 2020) libraries ggplot2, dplyr, viridis, tidyr, gganatogram, gridExtra, gtable. Sankey flow diagrams were produced using R (Team 2020) libraries ggplot2, dplyr, viridis, tidyr, gridExtra, gtable, cowplot, and networkD3. Kaplan-Meier plots for survival analysis were produced using the R library survminer (Kassambara 2020).

## Data access

### Hosting results and reproducibility

OncoMX datasets are stored and accompanied by a BCO (Simonyan et al. 2017) compliant with IEEE standards and publicly hosted at https://data.oncomx.org. GT dataset and modified forms of cancer datasets with glycobiology relevance can also be accessed from https://data.glygen.org. All scripts for analysis can be found at https://github.com/hmdinger/glycosyltransferases-in-cancer.

## Acknowledgements

The authors wish to acknowledge dissertation co-mentor Linda Werling, committee members and readers Nathan Edwards and Anelia Horvath, and examiners Katherine Chiappinelli and Mark Walderhaug; from the GW Institute for Biomedical Sciences, Marc Wittlif (former), Colleen Kennedy, Norman Lee, and Alison Hall; from the Department of Biochemistry and Molecular Medicine, Nichole Webster, Olga Reghay (former), and Rong Li; IT support and computing expertise: Robel Kahsay and Dacian Reece-Stremtan; OncoMX teammates and collaborators Frédéric Bastian, Daniel Crichton, Nikhita Gogate, Samir Gupta, Evan Holmes, Heather Kincaid, David Liu, Daniel Lyman, Marc Robinson-Rechavi, and Vijay Shanker;; Will York, Michael Tiemeyer, and Rahi Navelkar for glycobiology discussions; Jonny Torcivia for help with BioXpress and scripting; past labmates Yu Hu, Yu Fan, Quan Wan, Ting-Chia Chang, and Brian Fochtman for significant algorithm, pipeline, and data contributions; current labmates, especially Xiying Ding, Naila Gulzar, Jonathon Keeney, Janisha Patel, Stephanie Singleton, and Tianyi Wang.

## Supplemental Information

S1. GLY_000004 – curated enzyme listed hosted at data.glygen.org

S2. gts_all_annotations – table generated by automated steps of GT generation pipeline with evidence retrieved supporting each enzyme’s inclusion in the list

S3. curation_notes – table of hits from automated GT retrieval portions flagged for manual review and curation with corresponding curation notes and decisions (when applicable)

S4. curated_gts_appended – curated list from current (2020) release of GLY_000004 appended with functional annotations from lists generated by Hansen et al. and Joshi et al.

S5. cancer_do_summary - Cancers with differential expression data. Cancers listed include all cancers with differential expression data in at least one version. Cancers with lighter gray text were either excluded from downstream analysis in v2.0 (studies CESC, PAAD, READ, SARC, and UCEC) or altogether excluded from v4.0 (studies CESC, PAAD, and SARC) due to low patient count for paired samples. Shaded rows were combined according to the corresponding CDO slim disease mapping in the v4.0-tissue dataset. Colorectal cancer (DOID: 9256) includes data from COAD and READ studies; kidney cancer (DOID: 263) includes data from KICH, KIRC, and KIRP studies; lung cancer (DOID: 1324) includes data from LUAD and LUSC studies.

S6. BioXpress_DEGs_by_version - Differentially expressed gene counts at various fold change thresholds. S7. DEGTs_in_cancer – the subset of all differentially expressed GTs in cancer with reported *P* < 0.05

S8. DEGT counts by log2FC

S9. DEGT_cancer_counts – DEGTs summarized by count of cancers in which they are over- or under-expressed

S10. Similarity of DEGT trends across cancers – figure showing counts of DEGT trends plotted against log2FC thresholds

S11. DEGT trend similarity – figure showing distribution of similar or different expression patterns across multiple cancer types

S12. sample counts

S13.enrichment by cancer

S14. enrichment_by_family – table of findings with *P <* 0.05 from enrichment analysis of different GT families in each cancer type

S15. human_mouse_gt_orthologs – table human GT gene symbols and UniProt accessions, corresponding mouse gene symbols and UniProt accessions for those genes identified as 1:1 pairwise orthologs from OMA, and annotation of whether OMA mouse orthologs are included in the GlyGen mouse GT list

S16. similarity_score_matrix – matrix with all scoring possibilities and criterion applied to analysis of comparison of differential expression in cancer with healthy expression in human and mouse orthologous genes

S17. Similarity impact in male anatograms – figure showing impact score of all differentially expressed genes in cancer summed and averaged for males in available tissues

S18. high_value_mirna_pairs – table of high-value DEGTs and the miRNAs that target them with corresponding DE details for both

S19. cancer DEGTS from lit mining

S20. known_biomarker_overlap – subset of DEGTs overlapping with EDRN and FDA biomarkers S21. survival_summary – survival analysis results for all 36 DEGT-cancer pairs

S22. Similarity impact in male and female diagrams – figure showing the impact score of all differentially expressed genes in cancer both summed and averaged for males and females in available tissues.

