## Supplemental Figures for "Differential expression of glycosyltransferases identified through comprehensive pan-cancer analysis"

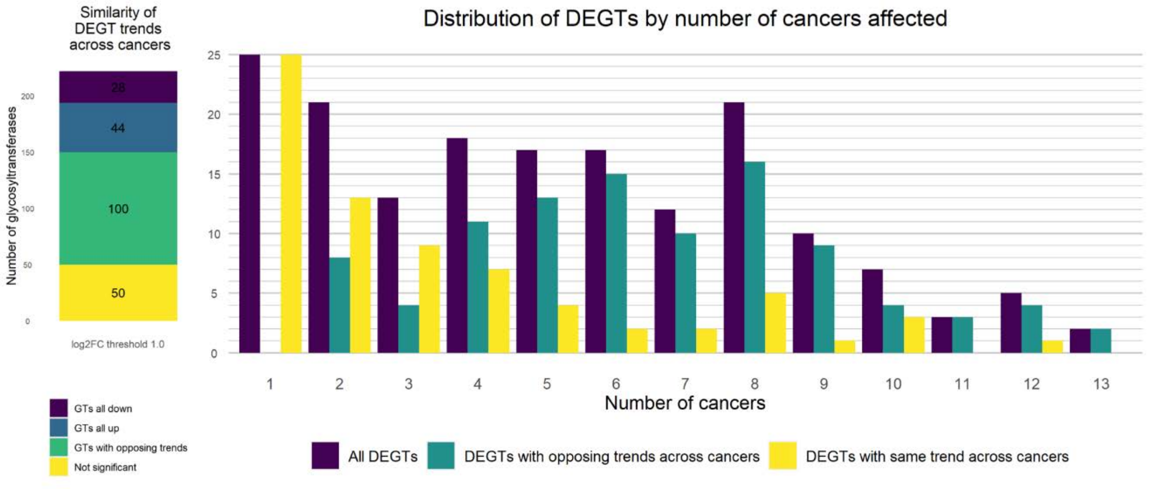

Similarity of DEGT trends across cancers
Number of glycosyltransferases
0
50
100
150
200
28
44
100
50
log2FC threshold 1.0
GTs all down
GTs all up
GTs with opposing trends
Not significant
Distribution of DEGTs by number of cancers affected
0
5
10
15
20
25
1
2
3
4
5
6
7
8
9
10
11
12
13
Number of cancers
All DEGTs
DEGTs with opposing trends across cancers
DEGTs with same trend across cancers

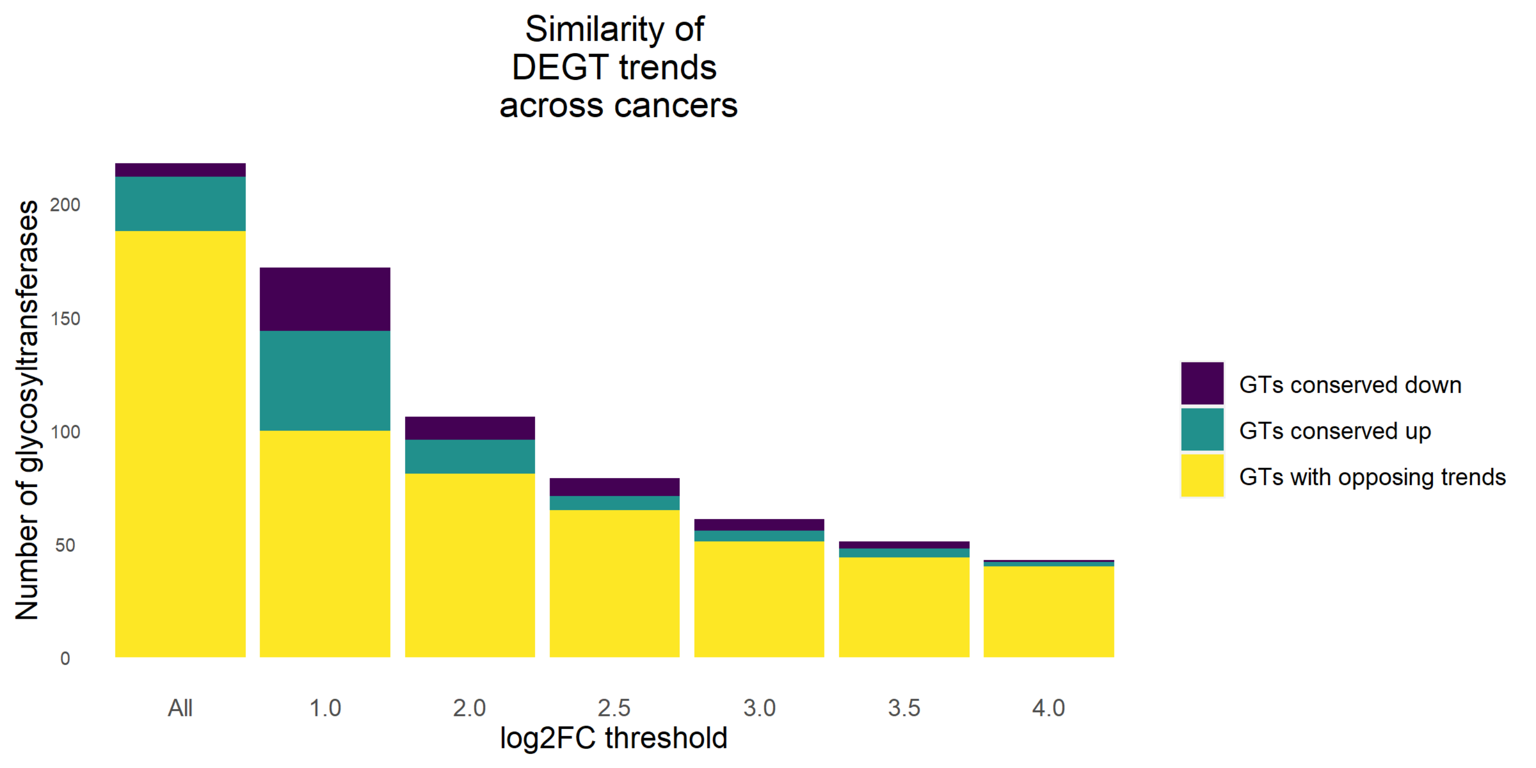

Similarity of
DEGT trends
across cancers
Number of glycosyltransferases
0
50
100
150
200
All
1.0
2.0
2.5
3.0
3.5
4.0
log2FC threshold
GTs conserved down
GTs conserved up
GTs with opposing trends

Impact of conservation on differential expression in cancer

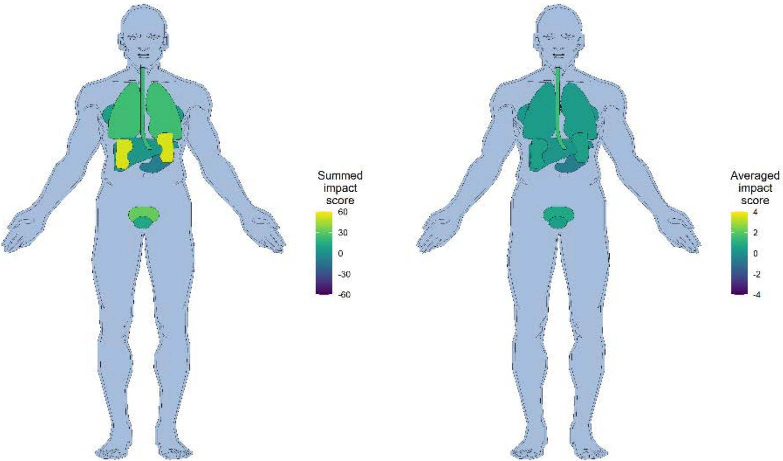


Summed for all genes

Averaged for all genes

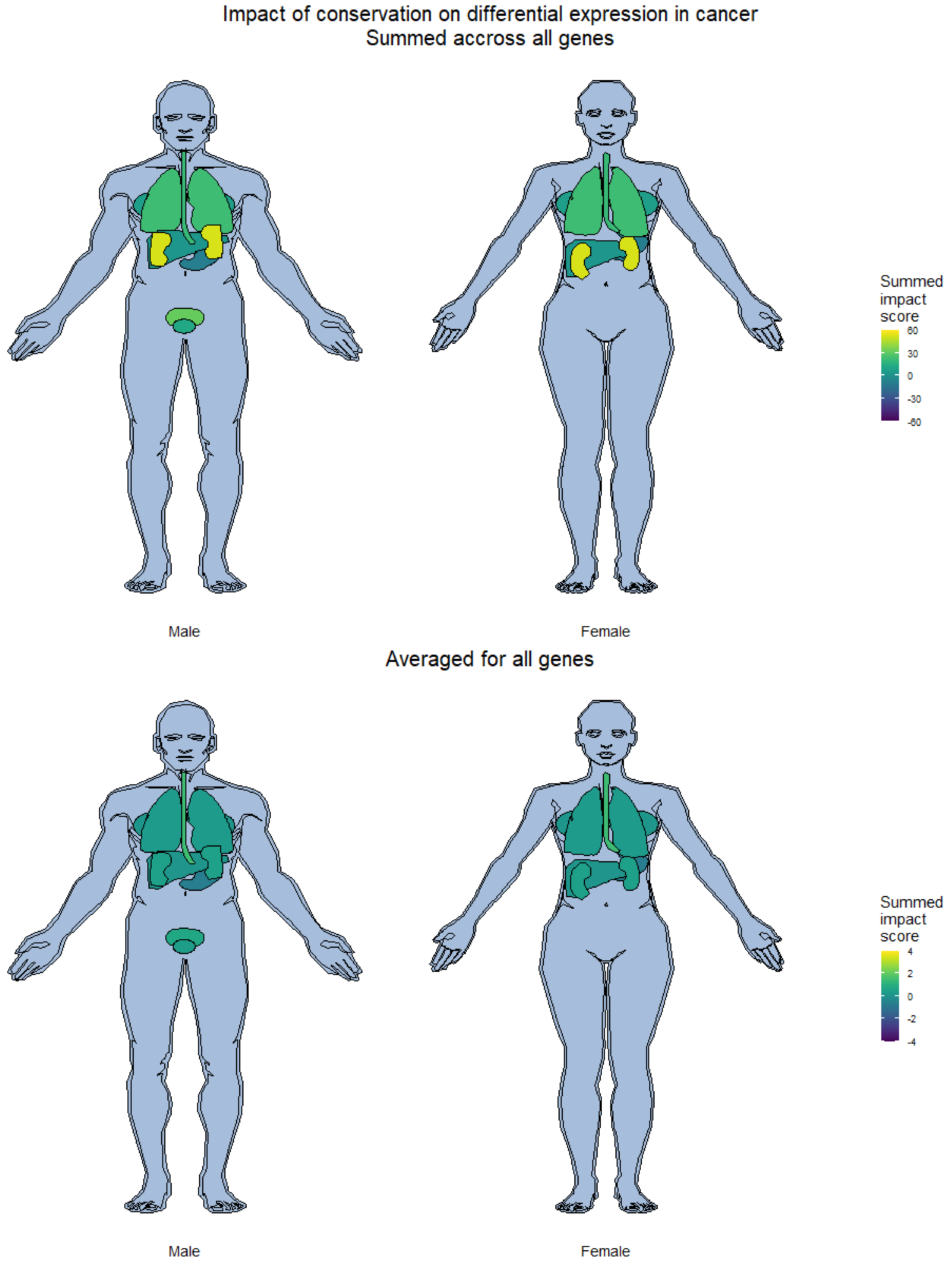
Impact of conservation on differential expression in cancer
Summed accross all genes
Summed impact score
60
30
0
-30
-60
Male
Female
Averaged for all genes
Summed impact score
4
2
0
-2
-4
Male
Female
